# Conformational Changes of RORγ During Response Element Recognition and Coregulator Engagement

**DOI:** 10.1101/2021.05.25.445650

**Authors:** Timothy S. Strutzenberg, Scott J. Novick, Ruben D. Garcia-Ordonez, Christelle Doebelin, Yuanjun He, Mi Ra Chang, Theodore M. Kamenecka, Patrick R. Griffin

## Abstract

The retinoic acid receptor-related orphan receptor γ (RORγ) is a ligand-dependent transcription factor of the nuclear receptor super family that underpins metabolic activity, immune function, and cancer progression. Despite being a valuable drug target in health and disease, our understanding of the ligand-dependent activities of RORγ is far from complete. Like most nuclear receptors, RORγ must recruit coregulatory protein to enact the RORγ target gene program. To date, a majority of structural studies have been focused exclusively on the RORγ ligand-binding domain and the ligand-dependent recruitment of small peptide segments of coregulators. Herein, we examine the ligand-dependent assembly of full length RORγ:coregulator complexes on cognate DNA response elements using structural proteomics and small angle x-ray scattering. The results from our studies suggest that RORγ becomes elongated upon DNA recognition, preventing long range interdomain crosstalk. We also determined that the DNA binding domain adopts a sequence-specific conformation, and that coregulatory proteins may be able to ‘sense’ the ligand- and DNA-bound status of RORγ. We propose a model where ligand-dependent coregulator recruitment may be influenced by the sequence of the DNA to which RORγ is bound. Overall, the efforts described herein will illuminate important aspects of full length RORγ and monomeric orphan nuclear receptor target gene regulation through DNA-dependent conformational changes.

**GRAPHICAL ABSTRACT:** 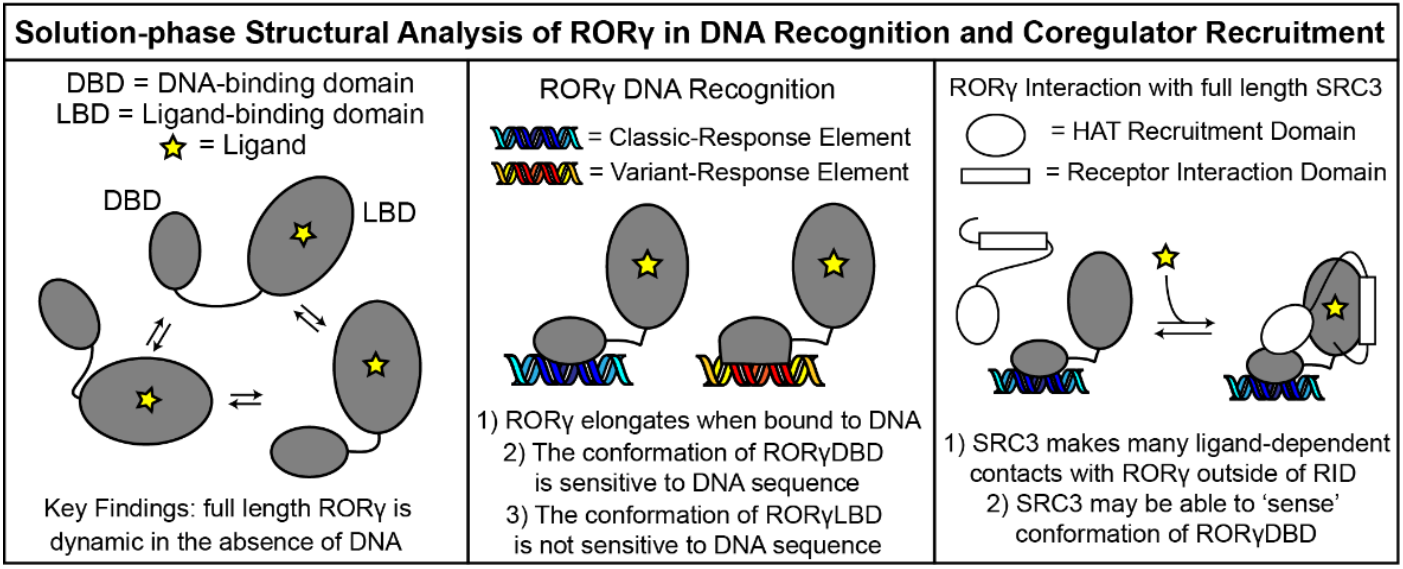

## INTRODUCTION

RORγ (gene name *RORC*) is a member of the nuclear receptor (NR) superfamily, acting as a ligand-dependent transcription factor to regulate biological pathways involved in metabolism, immune function, and cancer biology. RORγ isoform 1 (RORγ1) is widely expressed throughout the body [1, 2] and RORγ isoform 2 (RORγ2, RORγt) is expressed in lymphocytes. RORγ1 regulates gluconeogenesis in hepatocytes [3] and hypertrophy in adipocytes to reduce insulin sensitivity [4]. RORγ1 has also been found to play roles in driving progression of cancer, including triple negative breast cancer [5, 6], castration resistant prostate cancer [7], and pancreatic adenocarcinoma [8]. RORγ2 is the so-called ‘master regulator’ of IL-17 producing T helper cells (Th17) [9, 10]. Th17 cells play important roles in pathogen clearance (reviewed in [11, 12]) but can also drive pro-inflammatory autoimmune disorders (reviewed in [13]). Bi-allelic loss-of-function nonsense mutations in *RORC* have been observed in human patients whom present as immunocompromised [14]. RORγ2 also plays a critical role in thymopoesis where it promotes survival of double positive T cells [15, 16]. There has been significant effort towards development of RORγ antagonists/inverse agonists that inhibit RORγ activity to treat chronic autoimmune disorders. In rodents, pharmacological inhibition of RORγ phenocopies genetic depletion, producing resistance to autoimmune disorders [17-19] via inhibition of Th17 differentiation. Unfortunately, concomitant inhibition of thymopoesis also leads to the development of thymic aberrations and lymphoma [20-22]. Similarly, many clinical trials investigating oral administration of RORγ antagonists/inverse agonists for treatment of autoimmune disorders have been suspended due to safety concerns [23]. Development of a functionally selective modulator that blocks inflammatory Th17 pathways yet causes minimal disruption of thymopoesis could rescue RORγ as a viable drug target for autoimmune disorders.

Like most NRs, RORγ regulates the expression of target gene programs through a N-terminal DNA-binding domain (DBD) and a C-terminal ligand-binding domain (LBD). Unlike other nuclear receptors that can recruit coregulatory protein through a disordered N-terminal domain called activations function 1 (AF1), RORγ has a short N-terminus that does not have known activity. The RORγDBD recognizes specific DNA sequences called RORγ response elements (ROREs) via both major and minor groove contacts [2]. The RORγLBD recruits coregulator proteins with chromatin remodeling capabilities that promote transcription of RORγ target genes through activation function 2 (AF2). RORγ exhibits high basal activity which indicates that it recruits coactivators in the absence of exogenous ligands. RORγ is known to be responsive to cholesterol biosynthesis intermediates [24, 25], and oxysterols [26]. Further, mutations have been found that disrupt binding and activation by endogenous ligands but not synthetic ligands. These mutations result in loss-of-function that can be recovered by synthetic ligands which supports the model where the basal activity of RORγ is driven by endogenous ligands [27].

Structure-function analyses of RORγ have revealed important aspects regarding ligand-dependent transactivation of the receptor. However, these studies have been limited to analysis of the reconstituted LBD and do not explain several important functional observations that have been described in the literature. First, multiple groups have found that RORγ target genes can be driven through two unique RORE types termed classic- and variant-ROREs [5, 7, 28, 29]. Although both motifs share a common core [A/G]GGTCA nucleotide sequence the classic-ROREs have an A-rich 5’ flanking region while variant-ROREs lack the A-rich region. However, it is unclear if regulation of the RORγ target gene is sensitive to RORE type. Second, loss-of-function mutations have been found in regions of the receptor outside of the LBD including the DBD and the unstructured hinge region; for example mutations at the interface between the DBD and hinge region cause loss-of-function in Th17 polarization but not thymocyte survival [30]. Third, the *RORC* gene products are extensively regulated spatially and temporally after translation by translocation, post-translational modifications, and coregulatory protein interactions. Uncovering and defining these processes and mechanisms could potentially lead to new avenues that selectively manipulate the RORγ gene program based on RORE type by targeting regulatory pathways (reviewed elsewhere[31]).

To address these important questions, we applied a combination of exploratory data analysis of RORγ target gene programs and solution-phase structural analysis of full length RORγ. Analysis of published ChIP- and RNA-seq datasets suggests that classic- and variant-ROREs have different genomic distributions and that functionally distinct subsets of the RORγ gene program could be grouped by RORE type. Characterization of the structural basis of RORE recognition using structural proteomics revealed that full length RORγ2 elongates upon RORE recognition with an absence of long-range allosteric communication between the DBD and LBD. These data were combined with small angle x-ray scattering measurements to facilitate integrative modeling of the full length RORγ:RORE complex. To investigate coregulator engagement, we mapped ligand-dependent interactions with full length coactivator protein SRC3 using our structural proteomic platform. These studies also uncover that coregulators such as SRC3 may be able to ‘sense’ the RORE- and ligand-bound status of RORγ. Combining these observations, we propose a structural model where subsets of the RORγ gene program could be regulated in part through the response element sequence.

## RESULTS AND DISCUSSION

### DNA Recognition by Monomeric NRs

It is well documented that DNA sequence of the nuclear receptor response element can influence hetero- and homodimer NR activity, but this phenomenon is less understood for monomeric NRs such as RORγ. A key difference in monomeric receptors such as RORγ is the C-terminal extension (CTE) of the DBD that allows these receptors to distinguish between, and differentially bind to, structurally-similar response elements. DBDs from several monomeric receptors have been characterized and grouped into three classes depending on their response element sequence preferences and CTEs. For example; type I orphan receptor DBDs (including RORα and REV-ERBβ) prefer AANTAGGTCA[32, 33], type II DBDs (such as NGFI) prefer AAAGGTCA [34, 35], and type III DBDs (e.g. ERRβ and FTZ-F1) prefer CTAGGTCA [36, 37]. Interestingly, chromatin immunoprecipitation experiments have revealed that RORγ can recognize two classes of ROREs (see **FIGURE 1A**) called classic- and variant-ROREs. This indicates that RORγ can recognize and activate ROREs recognized by both type I and III DBDs. Some specific examples are that RORγt activates expression of lineage defining cytokine IL-17 in Th17 cells using a classic-RORE [38] while RORγ activates expression of the androgen receptor in prostate cancer using a variant-RORE [7].

**FIGURE 1:**
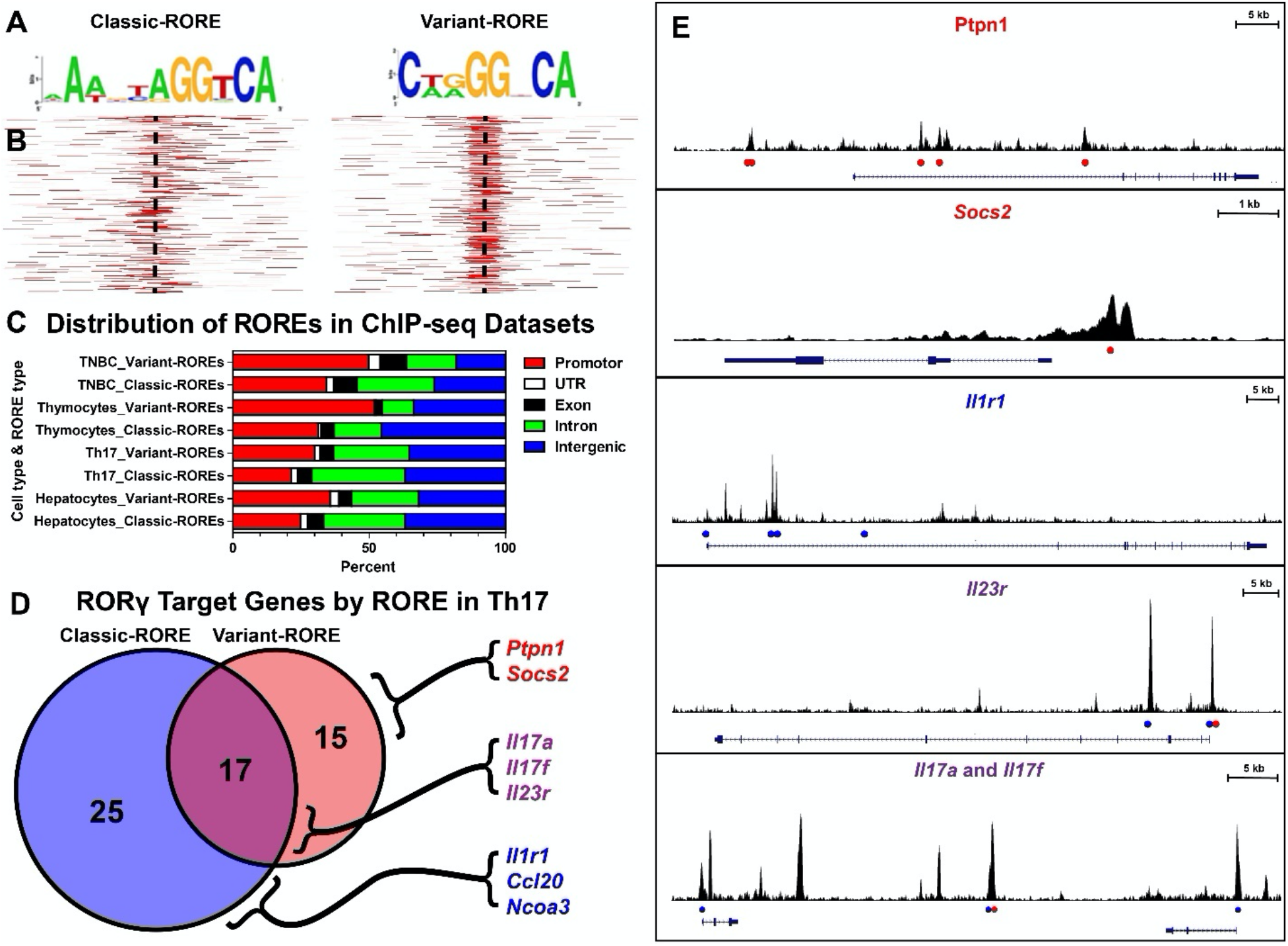
Genomic distributions of RORE and influence on RORγ target gene program. The motif logos for classic and variant ROREs are shown in **panel A**. Representative heatmaps of ChIP peaks containing classic- and variant-RORE are shown on the left and right respectively in **panel B**. Each row is a 10 kb promoter region centered on the TSS indicated by a dashed line. Red segments are 1 kb segments that contain a classic- or variant-RORE. An overview of global RORE annotation from disparate biological contexts is shown as a horizontal bar chart in **panel C**. Samples for triple negative breast cancer (TNBC) originated from the HCC70 cell line [5]. Naïve T cells were isolated form C57BL/6 and polarized toward Th17 *ex vivo* [28].

### RORγ response elements have distinct genomic distributions and regulate subsets of the RORγ target gene program

We sought to determine if classic-ROREs and variant-ROREs could be distinguished from each other with exploratory bioinformatic analyses. We explored validated ChIP-seq datasets from disparate biological contexts and found that classic- and variant-ROREs have distinct genomic distributions [5, 22, 28, 29]. Briefly, we determined qualitative attributes of classic- and variant-ROREs by annotating ChIP peaks by RORE type and characterized global genomic and promoter occupancy of RORγ from Th17 cells cultured *ex vivo*, thymocytes, hepatocytes, and the triple negative breast cancer (TNBC) cell line HCC70. It should be noted that ‘ROREs’ identified with this analysis are simply putative RORγ binding sequences and are not considered bonafide response elements until functionally validated. As shown in **FIGURE 1B**, heatmap analysis of promoter regions revealed that classic-ROREs tend to be distributed more frequently along gene loci compared to variant-ROREs that appear more localized to promoters and transcription start sites. Interestingly, this trend of classic-ROREs being found less in promoter regions and more in intronic (within introns) or intergenic regions (between genes) of the genome when compared to variant-ROREs was observed in every biological context examined (**FIGURE 1C**). Although these trends suggest that the RORE types are distinct, they do not provide a clear model for how RORγ target gene programs are regulated by the two ROREs.

Thymocytes and hepatocytes were isolated from C57BL/6 and C57BL/6J mice, respectively [22, 29]. RORγ target genes within Th17 cells cultured *ex vivo* were annotated based on an analysis of previously published RNA-seq and ChIP-seq datasets [28] and are shown in **panels D** and **E**. Genes identified by RORγ knockdown were first identified based on differential RNA-seq. The RORγ target genes were identified by cross referencing for ChIP peaks containing ROREs within loci that were reduced by RORγ knockdown. The target genes were then classified by the RORE type that may be regulating the expression of the loci and this is shown as a Venn diagram in **panel D**. Representative ChIP-peaks are mapped onto putative target gene loci in **panel E**. Target genes are labeled above the peaks and the scale of the genomic segment is shown on the top right inserts. The locations of classic- and variant-ROREs identified are indicated by blue and red circles, respectively.

To determine if RORγ target gene functions could be dissected by RORE-type, we analyzed previously published differential RNA-sequencing of Th17 cells cultured *ex vivo* [28]. We determined that 156 RNA transcripts showed statistically significant change upon knockdown of RORγ in these cells. We cross referenced the significantly altered gene transcripts with annotated ChIP peaks to identify 57 putative RORγ target genes. These included known RORγ target genes such as *Il17a, Il17f, Il23R*, and *Il1r1*. Gene set enrichment analysis (GSEA) showed significant enrichment of pathways related to T helper cell differentiation, Th17 activation, and IL-23 signaling, indicating that the analysis performed as anticipated **(SUP.FIG. 1A)**. We then separated the RORγ target genes based on the RORE type and found a somewhat even distribution of RORγ target genes regulated by classic-, variant-, or both RORE types, as shown in **FIGURE 1D**. The ROREs that putatively regulate the expression of these genes are mapped to their respective loci in **FIGURE 1E**. GSEA of RORγ target genes regulated by classic-ROREs showed enrichment of genes involved in Th17 cell activation and IL-23 signaling pathways, both of which are pro-inflammatory (**SUP.FIG. 1B**). Interestingly, the same analysis of variant-ROREs did not enrich Th17 cell activation pathways but rather showed significant enrichment of the JAK/STAT signaling pathway (**SUP.FIG. 1C**). Two genes contributed to the enrichment of this pathway, *Socs2* and *Ptpn1*, whose gene products act as protein tyrosine phosphatases that inhibit JAK/STAT signaling. JAK/STAT signaling is a major signaling paradigm throughout in the immune system (reviewed elsewhere[39]) and it is possible that RORγ could indirectly inhibit JAK/STAT signaling in Th17 cell differentiation through variant-but not classic-ROREs.

These observations warrant further exploration as it may be possible to regulate the RORγ target gene program based on RORE type. For instance, certain cytokine and signaling protein treatments have been shown to make Th17 cells more pathogenic (drive inflammatory disease) versus homeostatic (participate in pathogen clearance but not inflammatory disease) [40-42]. It may be possible that the signaling pathways that drive pathogenicity also emphasize aspects of the RORγ target gene program unequally. If cellular pathways exist to control RORγ activity on the basis of RORE type, then uncovering them and dissecting a mechanism, would unveil new avenues for developing functionally selective RORγ modulators that disrupt pathogenic, but not homeostatic activities of RORγ within Th17 cells.

### Structural analysis of full length RORγ

First we sought to characterize the structural basis of RORE recognition by RORγ using differential hydrogen-deuterium exchange coupled to mass spectrometry (HDX-MS). HDX-MS monitors conformational changes via altered backbone amide hydrogen exchange with deuterated solvent [43]. HDX-MS has been used previously to characterize long-range allostery in heterodimeric NRs such as PPARγ [44], VDR [45, 46] as well as monomeric NR LRH1 [47]. Initial digestion optimization of RORγ2 gave limited sequence coverage in the DBD. This was likely due to the high positive charge density, as basic residues cause missed cleavages by pepsin [48]. To optimize coverage of RORγ for HDX-MS analysis, we developed a method that improved peptide identifications from immobilized acid-stable proteases (described in Materials and Methods). Using this protocol, we were able to improve RORγ coverage from 77% to 98%, adding coverage to the DBD and allowing for the study of DBD-LBD crosstalk.

We first characterized how the DBD and hinge influence LBD dynamics by comparing the RORγLBD construct with the full length RORγ2 construct. Differential HDX-MS analysis revealed that the LBD within the context of full length RORγ2 was generally more protected from solvent exchange (and thus less dynamic) than the isolated RORγLBD (**FIGURE 2A**) with strong protection at the coactivator interaction surface (activation function 2, AF2) at helix 12 (H12) and the β-sheet region (BSR). Unexpectedly, strong protection was observed within H1 which connects the LBD to the hinge (**FIGURE 2F**). Overall, the LBD appears to be more stable within the context of the full-length protein likely due to hinge-LBD contacts as described below. It is unlikely that this observation is due to different ligands being bound during purification as the two proteins are both purified from *E. coli* in similar growth conditions.

**FIGURE 2:**
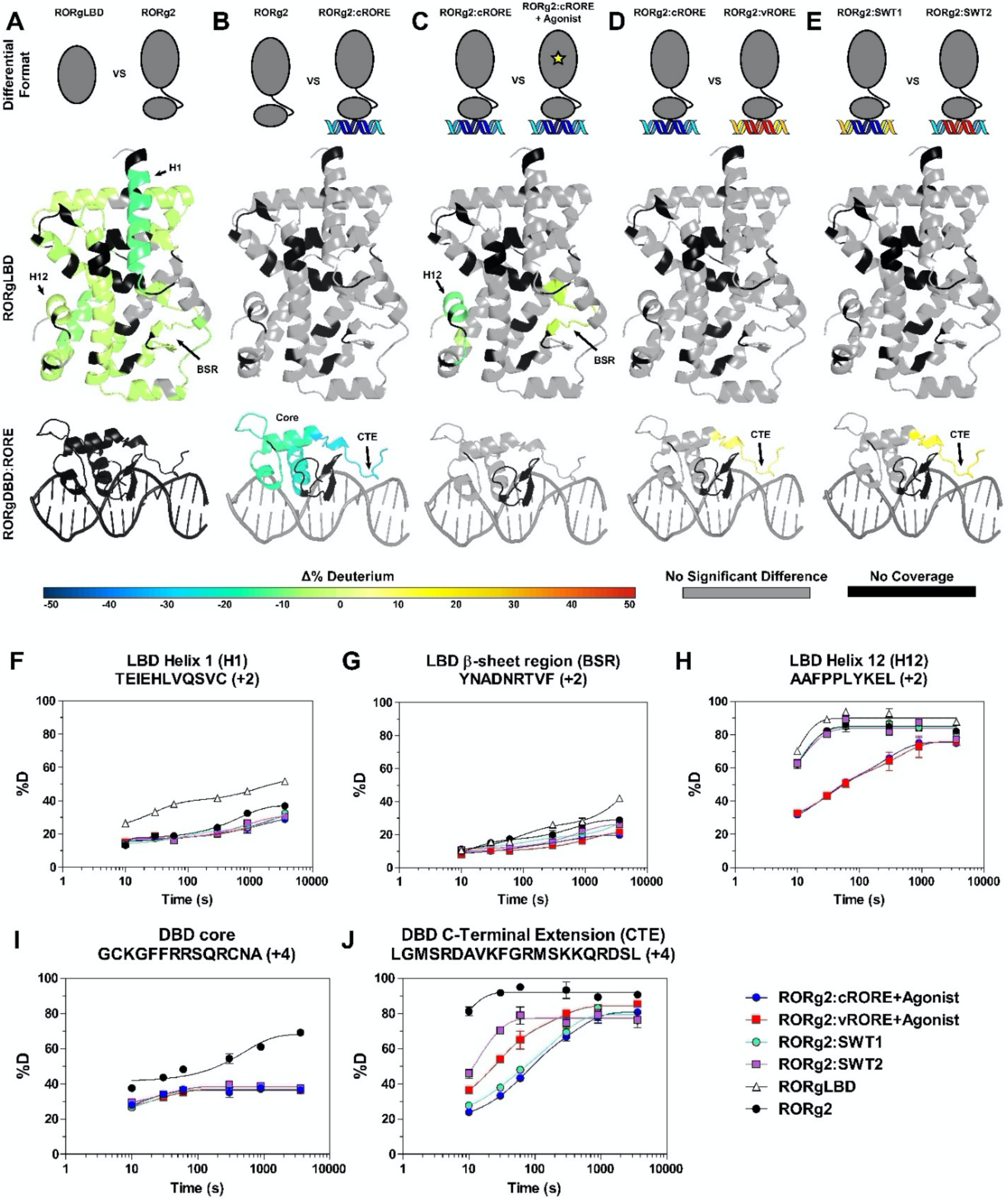
Characterization of RORγ bound to classic- and variant-ROREs by HDX-MS. An overview of differential HDX-MS experiments performed are shown in **panels A-E**. Each specific experiment is shown in each column and the cartoon on top depicts the two samples that are being compared. HDX-MS data is overlayed onto RORγLBD crystal structure (PDBID:3L0L) of and a RORγDBD homology model (templated from RevERα DBD PDBID:1HLZ) complexed with a synthetic RORE (modeled using 3D-DART [50]). Each color indicates change in percent deuterium incorporation (Δ%D) according to the color scale shown below. Representative deuterium build-up plots for peptides are shown in **panels F-J**. The legend for the deuterium incorporation plots is shown on the bottom right.

We next characterized changes in solvent exchange of RORγ binding to a conserved 30 bp sequence that regulates IL-17 expression through a classic-RORE [38, 49] (referred to hereafter as cRORE). Comparison of RORγ2 to RORγ2:cRORE revealed extensive protection to exchange in the DBD upon cRORE binding, with no changes observed in the LBD (**FIGURE 2B**). Analysis of deuterium build-up plots showed intermediate exchange kinetics of the DBD core consistent with pre-existing secondary structural elements in that region (**FIGURE 2I**). The same plots also show that protection from exchange occurs at both early and later time points suggesting further stabilization of these elements upon binding RORE. As shown in **FIGURE 2J**, the exchange kinetics of the C-terminal extension (CTE) were fast, consistent with disordered loop-like structural elements. We observed strong protection from exchange at the CTE at every timepoint in the presence of cRORE, suggesting this disordered region likely participate in engaging DNA. It is unclear whether these amides are protected by direct interaction with DNA or by the formation of secondary structure. HDX analysis revealed extensive stabilization of the DBD with no change in dynamics observed in the LBD. This suggests that there was no allosteric communication between the DBD and LBD in response to RORE recognition. Thus, it is unlikely that DNA binding impacts coregulator interactions mediated by the AF2 surface on the LBD.

Since RORγ purified from E. coli is in an inactive apo conformation, we next examined if the DBD dynamics are altered upon ligand binding by comparing the dynamics of RORγ2:cRORE ± full agonist SR19547. SR19547 had previously shown extensive protection to H12 which correlates with coactivator peptide affinity [27]. As shown in **FIGURE 2C**, HDX-MS analysis revealed strong protection to exchange in H12 and modest protection within the β-sheet region as expected, indicative of ligand binding and consistent with the binding pose. However, ligand binding did not alter exchange kinetics outside the LBD which is consistent with a lack of allosteric communication between the LBD and DBD. Importantly, the ligand binding signature appears to be dampened in the full-length protein likely because the LBD is stabilized in the context of the full-length protein likely driven by hinge-LBD contacts. However, we did not observe perturbation to exchange kinetics in these regions when we alter ligand-binding status. This could be due to the experimental time window used to monitor solvent exchange where we do not detect changes occurring at the very fast (<10 s) or on the very slow (>4 h) regimes.

### HDX-MS reveals that the RORγDBD conformation is RORE sequence-dependent

Next, we characterized structural dynamics of RORγ2 bound to a 30 bp segment of DNA that was shown to regulate the androgen receptor in prostate cancer through a variant-RORE (referred hereinafter as vRORE) [7]. As shown in **FIGURE 2D**, direct comparison of RORγ2:cRORE to RORγ2:vRORE revealed significant differences between the complexes within the DBD CTE, whereas no perturbation of exchange kinetics was observed within the DBD core or the LBD. The exchange kinetics of the RORγDBD core are nearly identical regardless of the RORE type (**FIGURE 2I**). To confirm that these observations are due to the sequence of the response elements and not the sequences of the 3’ and 5’ flanking regions, RORE switch oligos (called SWT1 and SWT2) were generated that swapped the 12 bp segment containing the RORE between the cRORE and vRORE oligos. Differential HDX-MS analysis of RORγ2:SWT1 and RORγ:SWT2 revealed nearly identical exchange kinetics as was observed between RORγ2:cRORE to RORγ2:vRORE (**FIGURE 2E**). These results indicate that the DBD core is equally stabilized by both ROREs and that the DBD CTE is more stabilized when bound to classic-ROREs. These analyses demonstrate that the dynamics of the DBD are sensitive to the specific nucleotide sequence of the RORE. Although this observation reveals a structural mechanism for sequence specific RORE recognition by RORγ, the lack of communication between the DBD and LBD/coactivator interaction surface suggests that ligand binding is uncoupled from this mechanism and that the DBD and LBD are uncoupled from each other.

### XL-MS analysis of RORγ RORE binding

We performed differential XL-MS analysis of RORγ2, RORγ2:cRORE, and RORγ2:vRORE and compared relative changes in crosslink abundance as a measure of side chain residency and reactivity using label free quantitation. In these experiments we identified and quantified 131 crosslinks that are represented in the heatmap shown in **FIGURE 3A**. The identified crosslink sites involved several regions of interest, including the DBD core, the C-terminal extension, multiple regions within the hinge, and multiple sites of the LBD. This analysis demonstrated that there are more RORγ2 crosslinks detected in the absence of DNA than in the RORγ2:vRORE and RORγ2:cRORE complexes. The identified crosslinks were placed into 5 groups based on a k-means clustering algorithm [51]. Clusters 1 and 2 contained crosslinks that were less abundant in samples treated with either vRORE or cRORE. Cluster 3 contains crosslinks that did not change in abundance when RORγ2 was bound to either RORE. Cluster 4 was a smaller group of crosslinks that increased in abundance in both vRORE and cRORE treatment groups. Interestingly, crosslinks in cluster 5 were not found in the absence of DNA or in the RORγ2:vROREcomplex, yet were abundant in the RORγ2:cRORE complex. To gain better insight into the conformational changes of RORγ upon RORE recognition, we mapped the crosslinks from each cluster to the primary sequence of RORγ2 (**FIGURE 3B**). Cluster 1 crosslinks that show a large decrease in abundance in presence of RORE, mapped to DBD-hinge and DBD-LBD interdomain crosslinks. The 2^nd^ cluster of crosslinks with somewhat lower abundance in the presence of ROREs mapped back to the CTE-hinge. There were also some AF2-hinge crosslinks in this cluster. The presence of interdomain crosslinks in RORγ2 in the absence of DNA indicates that the receptor samples conformations where the DBD is in close proximity to the LBD and portions of the hinge. To reconcile this with our HDX-MS results, we interpret this to mean that apo RORγ samples conformations that include DBD and LBD proximity that result in crosslink formation but do not alter LBD backbone dynamics in the ensemble. The decrease in interdomain crosslink abundance can be broadly interpreted to indicate that the DBD becomes more distant from the hinge and LBD and that RORγ elongates upon RORE recognition.

**Figure 3:**
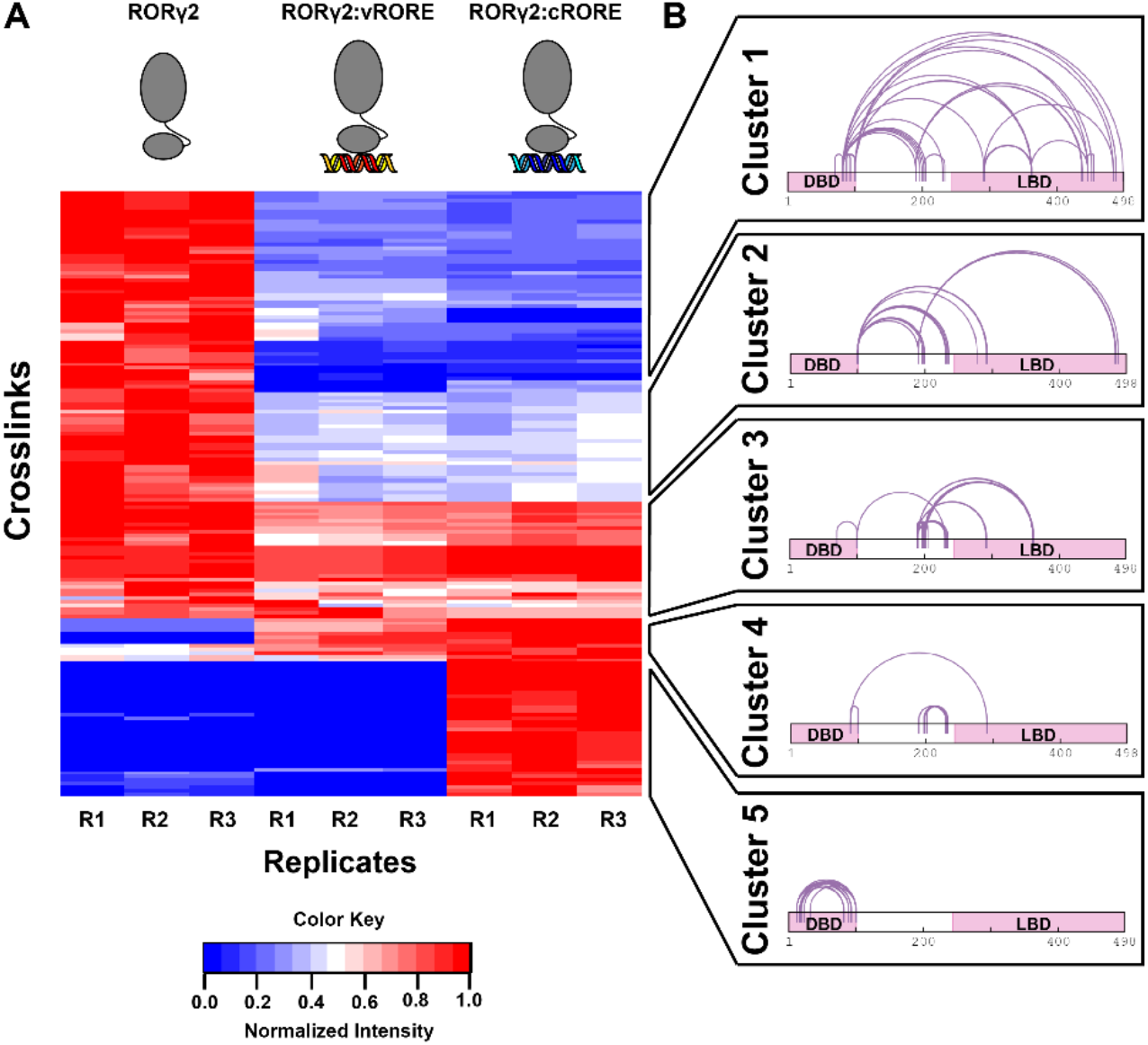
Conformational changes of RORγ2 upon RORE recognition determined by XL-MS. All quantifiable crosslinks are depicted in a heatmap in **panel A**. Each row of the heatmap represents a single crosslink species and each column represents a single replicate from RORγ2 (no RORE), RORγ2:vRORE, or RORγ2:cRORE as depicted on the top of the panel. The crosslinks were clustered into 5 groups using k-means clustering and the clustering results are shown in **panel B**. The groups of crosslinks were mapped to the primary sequence of RORγ2. The boundaries of the DBD and LBD are indicated by pink bars.

The 3^rd^ cluster of crosslinks that did not change substantially in all sample groups mapped to hinge-LBD crosslinks. This observation would suggest that regions of the hinge are in proximity of the LBD regardless of DNA-bound status. The smaller 4^th^ cluster consist mainly of hinge-hinge crosslinks. The changes in intra hinge-hinge crosslinks may be the consequence of rearrangement of RORγ upon RORE recognition in general, as they do differ between complexes with vRORE and cRORE. Interestingly, the 5^th^ cluster that is found exclusively in the RORγ2:cRORE complex is exclusively CTE-DBD crosslinks. These observations again highlight that the C-terminal extension can only crosslink with the DBD core when bound to a classic-RORE, suggesting that the C-terminal extension is proximal to the DBD core only when bound to cRORE but not vRORE. Overall, these findings are consistent with the HDX-MS data and suggest that the uncoupling of the RORγDBD and LBD is likely driven by an elongation that separates the DBD from the LBD.

### SAXS Illuminates Shape and Volume of RORγ:RORE Complexes

To further investigate if the conformational changes and domain-domain proximity observed by HDX- and XL-MS are consistent with an elongated multi-domain structure, we performed small angle X-ray scattering (SAXS) experiments using RORγ:RORE complexes. Unfortunately RORγ2 is not stable through freeze-thaw cycles, so we were unable to measure SAXS for the protein in the absence of RORE. The RORγ2:cRORE complex was monodispersed, and the scattering pattern was consistent with a lack of interparticle interactions or aggregation (**FIGURE 4A**). Analysis of the Guinier region (low q) showed that the radius of gyration and maximum distance were measured to be ∼55 Å and ∼200 nm, respectively. These measurements are comparable to other NR:DNA complexes reported previously [52]. As shown in **FIGURE 4B**, the pair distance distribution analysis shows a monomodal distribution with a rightward tail which indicates that the complex is more similar to a ‘rod-like’ shape than a globular sphere or ‘dumb-bell’ type shapes that would have a gaussian or bimodal pair distance distribution, respectively. Similar analyses of the RORγ2:vRORE complexes, resulted in data nearly superimposable to that obtained with the cRORE indicating unsubstantial change in the overall volume and shape of the different complexes. Importantly, these data support that there is no global conformational change of RORγ when bound to classic- or variant-ROREs.

**FIGURE 4:**
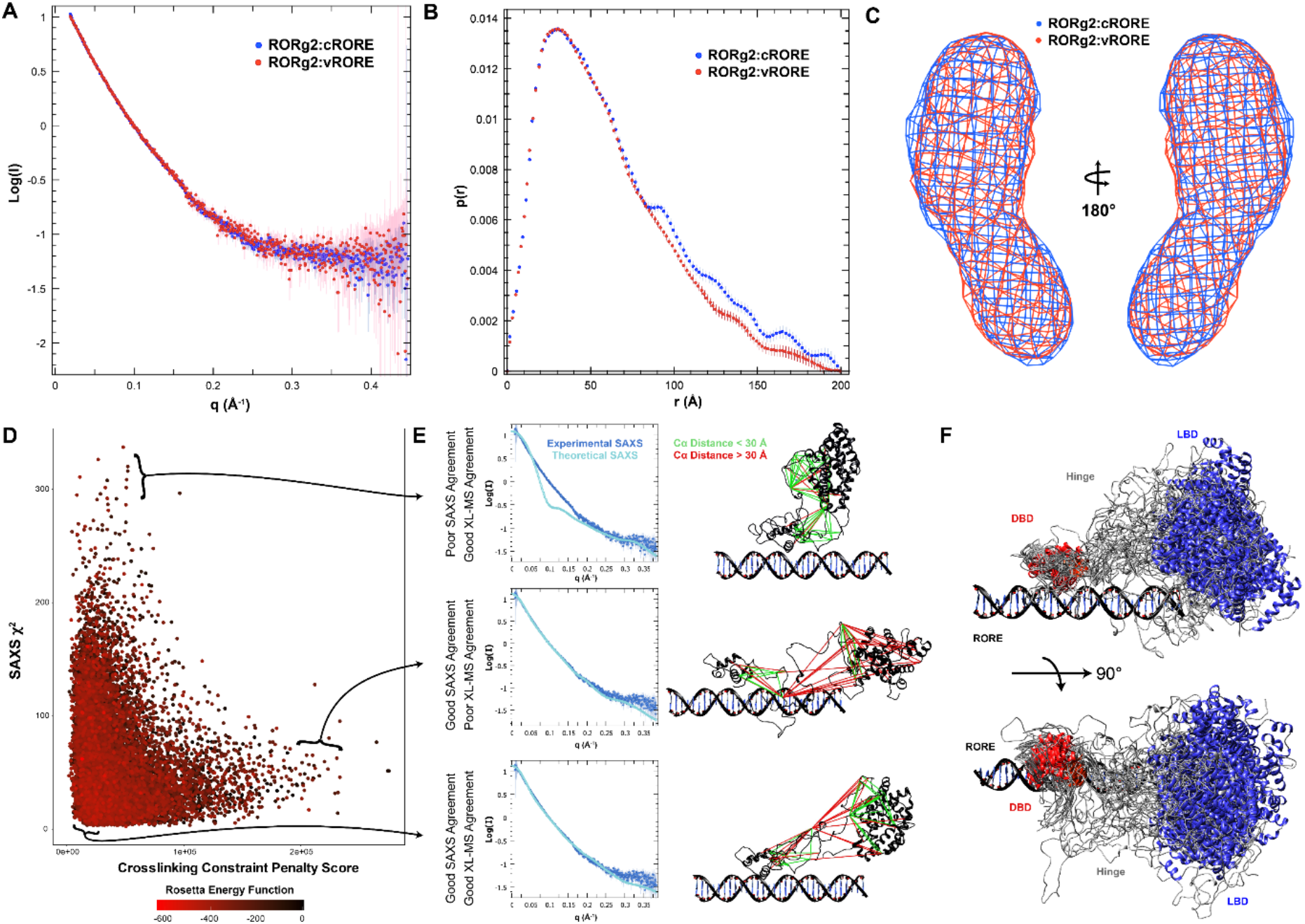
SAXS analyses of RORγ:RORE complexes. SAXS plots for RORγ2:cRORE and RORγ2:vRORE are shown in **panel A**. Pair distance distribution analysis is shown in **panel B**. Electron densities from *ab initio* modeling using DENSS are shown in **panel C**. The results from a modeling procedure using Rosetta are displayed as a dot plot in **panel D**. Each dot represents a structural model of RORγ2:RORE. Each structure is scored based on agreement with SAXS (SAXS χ^2^, Y-axis), XL-MS (Cα distance flat harmonic penalty, X-axis), and total Rosetta energy function score (colored coded according to legend at the bottom). Representative structures and data fittings are shown in **panel E**. Experimental and representative SAXS plots are shown on the left and representative atomic models and crosslink constraints are shown on the right. The 33 models that scored in the top 10% of both SAXS and XL-MS agreement are show in **panel F**.

To extract additional volume and dynamics estimates from the RORγ:RORE SAXS analyses, several computational analyses were performed. First, *ab initio* modeling using DENSS [53] generated model electron densities to create the pairwise distance distribution plot shown in **FIGURE 4B**. Over 50 electron densities were averaged into a single model presented in **FIGURE 4C**. Using the gold standard Fourier shell correlation approach [54], the resolution of the reconstructions is estimated to be ∼30 Å and there was unsubstantial qualitative differences between the density reconstructions between the RORγ2:cRORE and RORγ2:vRORE complexes. Interestingly, the electron density reconstructions appear to have two lobes. Second, we performed an ensemble modelling technique called EOM [55] that incorporates *a priori* information including structures and domain architecture of RORγ. Specifically, the RORγDBD:RORE and RORγLBD domains were treated as rigid bodies separated by a flexible hinge. Using this approach over 30,000 models were generated with a calculated theoretical SAXS plot for each structure. The theoretical SAXS curves were then fitted to the experimental SAXS plot where models that disagreed with the experimental data were removed. This approach yielded 4 structures that offer explanation of the observed SAXS plot and density reconstruction. To illustrate the results, the 4 structures were docked into the electron density reconstructions as shown in **SUP.FIG. 1A**. To demonstrate the diversity of the resulting ensemble of models, the structures were aligned to the RORγDBD:RORE as shown in **SUP.FIG. 1B**. In this view, the RORγLBD appears to sample several different orientations in space relative to the DBD:RORE. This approach highlights how the density might be better explained by multiple models rather than a single model. Third, to better understand how the XL-MS and SAXS results constrain the structure of RORγ2, we developed a Rosetta modeling pipeline that is shown in **SUP.FIG. 1C**. Briefly, the EOM analysis was repeated 12 times in total to generate 65 different LBD and DBD:RORE pairs that agree with the SAXS data. Using RosettaScripts, we remodeled the hinge region and relaxed the structures. The XL-MS results were incorporated as Cα distance flat harmonic penalties to the Rosetta energy function as previously described [56]. This process was repeated until 14,417 structures were generated. Each structure was then evaluated for agreement with the SAXS data using Crysol 3.0 [57] (reported as SAXS χ^2^). The results from this workflow are represented as a dot plot which is shown in **FIGURE 4D**. The results suggest that the experimental constraints guide RORγ models to two opposing extremes. The first general model suggests that RORγ can be more globular which, while satisfying the crosslinking constraints, does not agree with the SAXS data (**FIGURE 4E top**). The second general model suggests that RORγ can be elongated which agrees with the SAXS measurements, but violates most of the crosslinking Cα distance constraints (**FIGURE 4E middle**). Importantly, there were models that were adequate in explaining both sets of experimental constraints and those structures were generally in between both extremes and are shown in **FIGURE 4E bottom**. We then examined the top scoring models by filtering for structures that were in the top 10% of SAXS (SAXS χ^2^ < 13.2) and crosslink (crosslink constraint penalty < 14,800) agreement (**SUP.FIG. 2D**). This approach resulted in 22 structures that are shown in **FIGURE 4F**. Overall, this integrated modeling analysis suggests that the LBD becomes distant from the DBD upon RORE recognition and that experimental measurements constrain the hinge and LBD in proximity of the 5’ end of the DNA response element. This model does explain why conformational changes in the DBD do not influence the dynamics of the LBD, as there is no direct contact, and the regions separating these domains is mostly disordered and not capable of transmitting DNA-bound status allosterically.

### Reconstituting full length RORγ:coactivator complexes on RORγ response elements

Although the results presented so far suggest that RORγ recognition of ROREs is not directly regulated by ligand-binding, indirect regulation mechanisms could exist through ligand-dependent coregulator interactions. RORγ has been shown to recruit members of the p160 steroid receptor coactivator (SRC) family and histone acetyltransferases to facilitate transactivation of its target gene program and this process is thought to require RORγ to be bound to a ligand. Specifically, RORγ has been found to interact with SRC1 [58, 59], GRIP/SRC2 [18], SRC3 [60], and P300 [61]. Interestingly, there is anecdotal evidence that different SRCs drive subsets of RORγ target genes. Specifically, it has been suggested that RORγt:SRC3 (*NCOA3*) interactions drive ‘pathogenic’ Th17 responses [62], whereas RORγt:SRC1 interactions drive thymopoesis and physiological Th17 responses [59, 60]. Interestingly, to date, RORγ-SRC interactions have only been explored structurally using small peptides derived from coregulatory protein NR interaction motif (NR-box) or with small receptor interaction domains (RIDs) from the large coregulatory proteins. To further expand on these studies presented here, we characterized the ligand-dependent interactions of RORγ2:cRORE with the full-length coactivator SRC3 using HDX- and XL-MS.

### Mapping SRC3 interaction surface on RORγ with HDX-MS

RORγ2:cRORE was reconstituted and treated with synthetic agonist SR19547 (5-fold molar excess) followed by incubation with recombinant full length SRC3 (1.5-fold molar excess) or protein buffer control and the complexes were analyzed by differential HDX-MS. As shown in **FIGURE 5A**, substantial perturbation in solvent exchange was observed only in the RORγLBD. As expected there was strong protection to solvent exchange in H12 (**FIGURE 5C**) at all timepoints suggesting stabilization of the secondary structural elements in this region. Significant protection to solvent exchange was also observed within H1’, suggesting additional SRC3 contact sites on the RORγLBD (**FIGURE 5B**). This protection was observed at every timepoint; however, the slope of the deuterium build-up curve was unchanged. This indicates that amides that were not previously involved in hydrogen bonds (disordered) became ordered leading to stable secondary structure.

**FIGURE 5:**
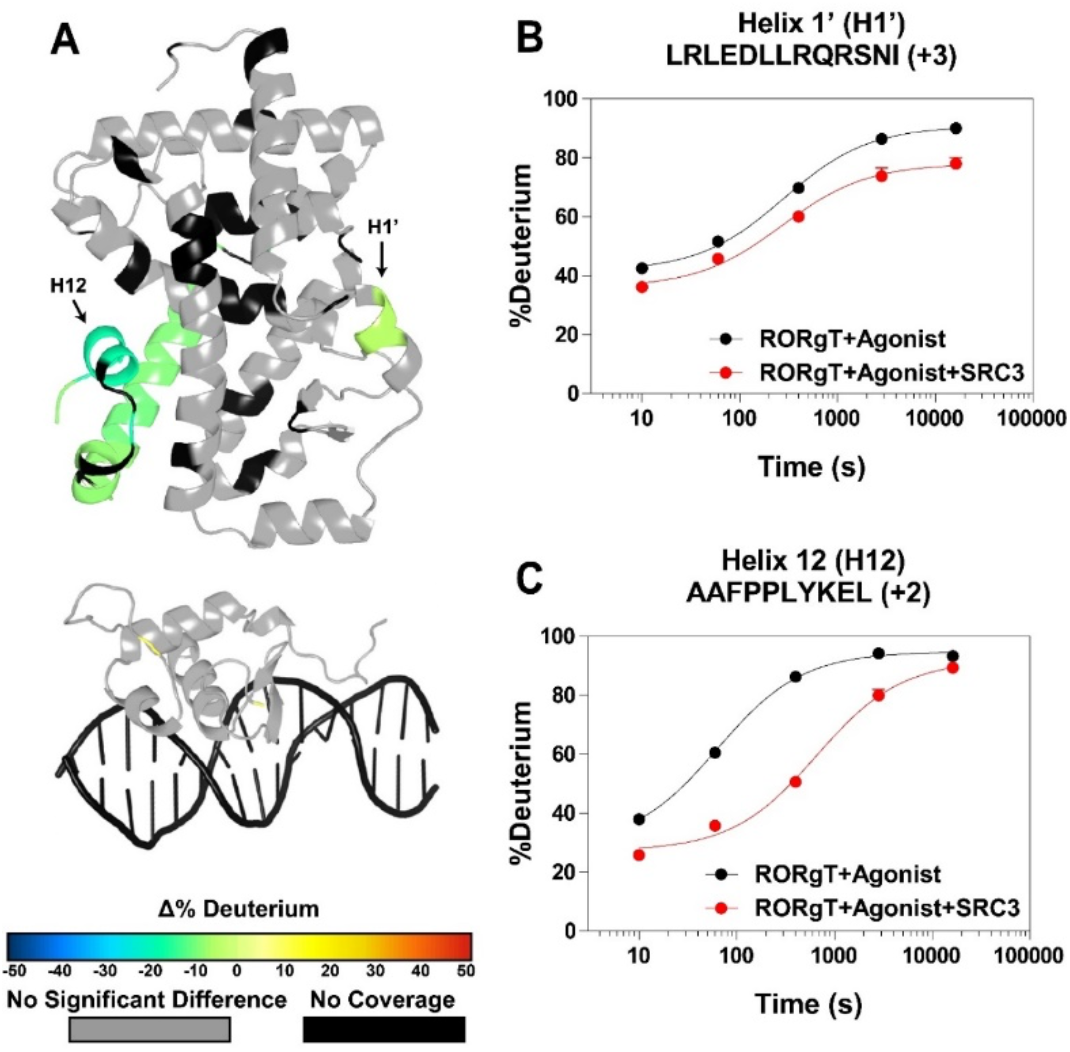
Characterization of full-length SRC3 interactions with RORγ:RORE complex by HDX-MS. The differential HDX of RORγ2:cRORE:SR19547 ± SRC3 was measured at 5 timepoints ranging from 10 s to 4 h. The Overlay of HDX data onto crystal structure(3L0L) of RORγLBD (top) and homology structure (template PDB: 1HLZ) for RORγDBD (middle) according to scale shown below in **panel A**. Representative deuterium build-up plots for peptides spanning helix 1’ are shown in **panel B**. Representative deuterium build-up plots for peptides covering helix 12 are shown in **panel C**.

### Mapping ligand-dependent RORγ-SRC3 contacts with differential XL-MS

RORγ2:cRORE was incubated with SRC3 ± SR19547. The samples were crosslinked with DSSO and subject to MS analysis. This resulted in the identification and quantification of 141 crosslinks including 117 intra- (SRC3-SRC3 and RORγ2-RORγ2) and 24 inter-protein (RORγ2-SRC3) crosslinks. The label-free quantitation results from this analysis are shown as a volcano plot in **FIGURE 6A**. Overall, we observed that the abundance of intra-protein crosslinks did not change between SR19547- and DMSO-treated samples (1 > log_2_ fold change > -1), and that inter-protein crosslinks were enriched in SR19547-treated samples (log_2_ fold change > 1). These results indicate that several regions of SRC3 have greater resident occupancy with RORγ when RORγ is bound to a ligand, an observation consistent with ligand-dependent RORγ:SRC3 interaction. To further explore the contact surface for this protein-protein interaction, we mapped the crosslinks to the primary sequence of RORγ and SRC3 based on fold change values. Crosslinks of similar abundances in both DMSO only and SR19547-treated samples are shown in **FIGURE 6B**. These crosslinks mostly consisted of inter-protein crosslinks. The crosslinks enriched in the SR19547-treated samples consisted of exclusively inter-protein crosslinks (**FIGURE 6C**).

**FIGURE 6:**
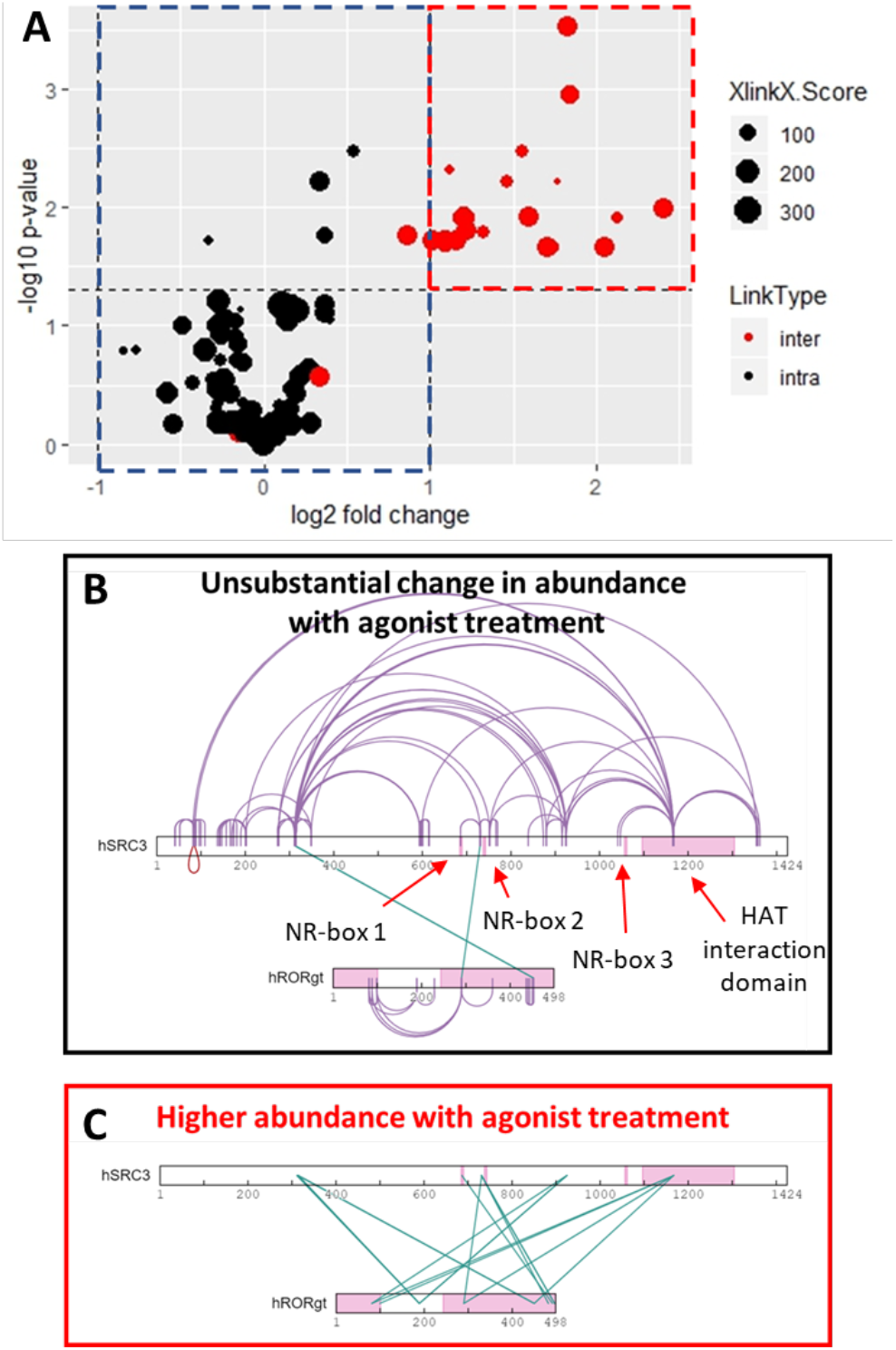
Mapping ligand-dependent SRC3-RORγ2 contacts by differential XL-MS. RORγ2:cRORE was incubated with SRC3, treated with agonist SR19547 or DMSO only, and crosslinked. The crosslink abundance was determined using label-free quantitation and the results comparting the agonist and vehicle treatment groups is shown as a volcano plot in **panel A**. Crosslinks are represented as dots where the area corresponds to the XlinkX score (the confidence of the identification). The crosslinks are colored black and red if they are intra- or inter-protein crosslinks, respectively. The crosslinks whose abundance did not change substantially with agonist treatment are mapped to the primary structure of SRC3 and RORγt in **panel B**. The crosslinks whose abundance were significantly higher in the agonist treated group were mapped back to the primary sequence of RORγt and SRC3 in **panel C**. The boundaries of the RORγ2 DBD and LBD are indicated by pink bars. The NR-box LXXLL motifs of SRC3 and the C-terminal histone acetyl transferase (HAT) interaction domain are indicated by pink bars.

This analysis and visualization revealed interesting and important aspects of the RORγ2:SRC3 interaction. Unexpectedly, SRC3 makes contacts in both the RORγDBD and LBD. More surprisingly, the HAT interaction domain of SRC3 makes contacts with the CTE of the RORγDBD. Recruitment of HATs to this domain of SRC3 is essential for the protein to influence chromatin structure. While HDX-MS analysis demonstrated that SRC3 interactions do not stabilize the backbone dynamics of DBD CTE, XL-MS reveals that the HAT interaction domain of SRC3 is in proximity of the DBD of RORγ. This would suggest that coactivator binding to RORγ may directly influence DNA modifications. Second, the first and second NR-box motifs of SRC3 contact RORγ2. This indicates that RORγ may have a preference for interacting with these motifs rather than the third NR-box motif.

## CONCLUSION

Structural analysis of intact NR:coregulator complexes has been notoriously difficult as there is a tradeoff between obtaining high resolution and capturing dynamics. While high resolution x-ray co-crystal structures of several NR heterodimers on response elements have been reported [63-65], obtaining these structures relied on heavily engineered protein constructs, and although these studies have generated several testable hypotheses, there is controversy regarding their biological relevance. Most often, structures solved with crystallography often fail to explain all observations made during solution-phase analyses. For example, in contrast with the PPARγ:RXR:PPRE crystal structure that suggested a compacted shape, solution-phase SAXS experiments suggested a more dynamic complex [52]. Further, the ligand-dependent recruitment of coactivators to the PPARγ:RXR heterodimer is positively cooperative with DNA binding [44]. Meanwhile, crystallization of NR:coregulators has involved the reduction of coregulators to short peptide fragments, preventing quaternary structure determination of NR:coregulator complexes. Fortunately, structural proteomics has exhibited high potential to advance studies of NR complexes [47].

Characterization of NR-coregulator interactions using single particle cryogenic electron microscopy (cryo-EM) has been sufficient to provide some structural insight to cooperation between DNA binding and coactivator recruitment; however, there remains an inverse correlation between dynamics and resolution. For example, while the VDR:RXR heterodimer is contains fewer disordered than most NRs, and solution-phase experiments demonstrate that the DBDs are capable of allosteric communication with the coactivator interaction surface of the LBD [45, 46], cryo-EM produced structures with a resolution of 10-12 Å, insufficient for precise determination of the mechanism of allostery [65]. Cryo-electron microscopy studies have also been attempted for AR:SRC3:P300 and ER:SRC3:P300, but the inherent disorder and dynamics of these systems has greatly limited resolution to around 30-40 Å [66, 67]. At this resolution, the general shape of the complex can be determined, but specific contact sites cannot be assigned to regions based on the density. Structural proteomic techniques such as HDX- and XL-MS can complement these efforts as they can precisely characterize interactions between proteins in the solution phase even when overall 3D structural information is limited or absent. While the HDX- and XL-MS platform provides no inherent atomic structural information, observations of intra-protein crosslinks infer that the amino acids involved in the crosslink must at some point come within ∼30 Å to form the crosslinking reaction transition state. Thus, our platform not only allows more precise analysis of protein-protein interactions but can also serve as a valuable complement to existing structure determination techniques by providing information about the solution-phase structural dynamics of large macromolecular complexes. For example, HDX-MS analysis revealed two contact sites in the RORγLBD that stabilize backbone dynamics and thus drive formation of the NR:coregulator complex whereas XL-MS revealed nine additional contacts sites between SRC3 and RORγ2 to complement the HDX-MS findings. As expected, we observed crosslinking of NR-box motifs 1 and 2 to the RORγ AF2 surface, likely driving the stabilization observed by HDX-MS in H12. This information would be difficult to ascertain from low resolution structural analyses alone.

The results presented here provide important context to full length monomeric NRs structure and function. Using similar techniques, Seacrist *et al*. found that monomeric NR LRH1 adopted a more globular complex on DNA such that the LRH1DBD transmitted information to the LRH1LBD [47]. In contrast, our studies suggest that RORγ becomes elongated upon DNA recognition effectively uncoupling the two domains. Regardless, it is still possible that the RORγLBD regulates the RORγDBD activity indirectly through coregulators. Previous studies further support this as it has been shown that the RORγDBD CTE is acetylated by P300 and this activity correlates with reduced DNA binding affinty [68]. Our XL-MS results show that SRC3 makes multiple ligand-dependent contacts with RORγ2:cRORE with a specific contact between the HAT domain of SRC3 and the CTE of RORγ. This finding is intriguing as it opens the possibility that SRC3 may interact with RORγ in a RORE-dependent manner or that DNA binding may influence coregulator association with HATs. Given that our structural proteomic approach is generalizable, it can be broadly applied to many different RORγ:coregulator complexes to explore ligand- and RORE-dependent regulation of RORγ, which is a focus of future studies.

## MATERIALS/METHODS

### Exploratory analysis of Classic- and Variant-ROREs

Data for the corresponding sequencing experiments were downloaded from the SRA database. The sequence adapters were trimmed and aligned to the respective genome (either mm9 or hg38) using bowtie2 [69]. Peak calling was done using the MACS2 algorithm [70] the peaks containing classic-(AAATAGGTCA) or variant-(BBCTAGGTCA [B indicates T, G, or C]) RORES (depicted in **FIGURE 1A**) were annotated using HOMER[71]. Qualitative analyses of RORE containing peaks was done using ChIPseeker [72] implemented in R Studio (version 1.3.1073). To determine differentially expressed genes in *RORC* knockdown Th17 cells, RNA-seq datasets quasi-aligned and quantified using salmon [73] and statistical analyses of the quantified read counts was done using DESeq2 [74] implemented in R Studio. The genes that were significantly different were then discarded if there was no RORE containing peak within 10 kbp of the transcription start site. We performed pathway analysis using Ingenuity Pathway Analysis software (QIAGEN).

### Materials

Unless otherwise stated, chemicals were purchased from Sigma-Aldrich (St. Louis, MO). SDS-PAGE analyses were done using 4-12% AnyKD™ gels (Biorad) according to the manufacturer’s protocol. Yeast extract, HEPES base, and HEPES free acid were purchased from Research Products Incorporated. Dioxane free isopropyl-β-thiopyranoside (IPTG) was purchased from Biosynth. Dithiothreitol was purchased from Fisher Scientific. Agar was purchased from Invivogen and prepared according to manufacturer’s instructions. Terrific broth (12 g tryptone [Fluka], 24 g yeast extract [Research Products Incorporated], 4 mL glycerol [Fisher Scientific], 9.4 g K_2_HPO_4_, 2.2 g KH_2_PO_4_ per liter of media) was prepared in house.

### Sample Preparation

#### RORγ Cloning and Expression

RORγ2 was codon optimized (Genewiz) and subcloned into the pESUMO vector using XhoI and XbaI. The RORγ1 construct was generated by PCR insertion with the Q5 mutagenesis kit (New England Biolabs). Both constructs were transformed in BL21 (DE3) expression *E. coli* and selected on Luria broth agar plates containing 100 ug/mL ampicillin. 6-8 colonies were picked for primary culture in 250 mL of terrific broth and cultured for 16 h at 37°C. 15 mL of primary culture was diluted into 1 L of terrific broth supplemented with 30 μM ZnCl_2_ and 50 ug/mL carbenicillin. The secondary culture was incubated at 37°C with shaking at 200 RPM until the optical density at 610 nm reached 0.5 after which the temperature was dropped to 16°C. protein expression was induced by addition of IPTG to 250 μM and the culture was incubated at 16°C for 8 h and then 4°C until harvest. The cells were harvested by centrifugation (4000 rcf for 10 minutes at 4°C), resuspended in ice cold NiNTA buffer 1 (50 mM HEPES pH 8.0 at 4°C, 500 mM NaCl, 10% glycerol, 25 mM imidazole) supplemented with 1X SigmaFast protease inhibitor cocktail, and pelleted again by centrifugation (4000 RCF for 10 minutes at 4°C). Harvested cells were flash frozen in liquid nitrogen and stored at -80°C until purification.

#### HisSUMO-RORγ Purification

RORγ constructs have poor stability through freeze-thaw cycles without the HisSUMO solubility tag or without being bound to DNA. To accommodate this, RORγ1 and RORγ2 constructs were purified in two stages. The first stage was to isolate highly pure HisSUMO-RORγ by affinity purification, and the second stage was RORγ:RORE complex formation followed by tag cleavage and removal. Harvested cell pellets were resuspended in ice cold NiNTA buffer 1 supplemented with 1X SigmaFast protease inhibitor cocktail, DNase, and lysozyme in a ratio of 4 mL of buffer per gram of cell pellet. The cells were lysed using a microfluidizer operating at 15,000 PSI and cooled to 4°C. The lysate was clarified by centrifugation (30,000 RCF for 30 minutes at 4°C) and the resulting supernatant was filtered with 0.45 μM cellulose vacuum funnel (Millipore). NiNTA resin (Qiagen) pre-equilibrated with NiNTA buffer 1 was added to the filtered supernatant (1 mL column volume per 40 mL supernatant) and the slurry was incubated at 4°C for 30 minutes with gentle rotation. The slurry was added to a flex column and the flow through was discarded. The column was washed 3 times with 10 column volumes of NiNTA buffer 1. The eluate was collected after adding 3 column volumes of NiNTA buffer 2 (50 mM HEPES pH 8.0, 500 mM NaCl, 10% glycerol, 250 mM imidazole) and incubating the slurry for 10 minutes. The elution was repeated twice, and the 3 eluates were pooled. The NiNTA eluate was slowly diluted 10X with ion exchange buffer 1 (25 mM HEPES pH 8.0, 100 mM NaCl, 1 mM TCEP, 5% glycerol). The protein solution was passed over a 5 mL HiTRAP heparin sulfate pre-equilibrated with ion exchange buffer 1 using an AKTA Pure™ fast protein liquid chromatography system sample pump operating at 5 mL/min. The column was washed with 5 column volumes of ion exchange buffer 1 using a gradient pump operating at 5 mL/min. The protein was eluted in a linear gradient of ion exchange buffer 2 (25 mM HEPES pH 8.0 at 4°C, 1000 mM NaCl, 1 mM TCEP, 5% glycerol) over 30 column volumes where 2 mL fractions were collected in glass vials. HisSUMO-RORγ elutes over 4 column volumes at ∼50% ion exchange buffer 2. After ion exchange, fractions containing HisSUMO-RORγ were pooled and concentrated until the protein concentration was ∼ 2 mg/mL (∼30 μM) based on absorbance at 280 nm with an extinction coefficient of 36330 cm^-1^M^-1^ (both RORγ1 and RORγ2 only have 2 tryptophan residues). The sample was split into 350 μL aliquots, flash frozen, and stored at -80°C until further use. The typical yield of this preparation is ∼ 4-6 mg of highly pure HisSUMO-RORγ per liter of expression media. Samples were taken throughout the purification process and analyzed by SDS-PAGE to assess purity.

#### RORγ:RORE complex purification

Synthetic dsDNA oligos were designed based on sequences of well-characterized and functionally relevant RORγ response elements. The sense and anti-sense oligos were synthesized, annealed, and HPLC purified (Integrated DNA Technologies). The lyophilized DNA was resuspended in molecular biology grade water to a concentration of 150 μM based on absorbance at 260 nm. To form RORγ:RORE complexes, 100 μL of RORE was added to 33 μL of 4X HBS (200 mM HEPES pH 7.5, 2M NaCl). The 30 μM HisSUMO-RORγ samples were thawed at room temperature, mixed, centrifugated (20,000 RCF for 3 minutes at 20°C) and added to the salt normalized RORE solution where the final concentration of RORE was an approximate 1.5 molar excess to RORγ. HisSUMO-RORγ:RORE complexes were formed over a 30 minute incubation at room temperature (RT). We confirmed protein-DNA complex formation by DNA retardation gel electrophoresis and analytical size exclusion chromatography. The HisSUMO solubility tag was cleaved enzymatically by addition of 10 μL of 20 μM His-tagged SUMO protease and incubating for 10 minutes at RT. RORγ:RORE solutions were purified, and buffer exchanged using size exclusion chromatography using an AKTA Pure™ system operating at RT. The solutions were manually injected onto a Superdex 200 10/300 (General Electric) pre-equilibrated with HEPES buffered saline (25 mM HEPES pH 7.5 at RT, 150 mM NaCl, 50 μM TCEP, 2% glycerol), eluted with an isocratic pump operating at 1 mL/min, and 200 μL fractions were collected in polypropylene 96 well plates. Fractions containing the RORγ:RORE of interest were pooled (typically 1.6 mL) and kept on ice. The yield was determined based on UV-Vis spectroscopy where the extinction coefficient of the DNA at 260 nm was used to determine concentration. The total complex recovery estimates typically ranged from 70-80%. To prepare the samples for XL- and HDX-MS, the samples were concentrated to ∼10 μM using 0.5 mL Amicon Ultra™ spin filtration devices with 50 kDa molecular weight cut off. To prepare the samples for HT-SAXS, dithiothreitol was added to 5 mM and the sample was concentrated to ∼1.0-2.0 mg/mL (30-40 μM) using a separate spin filtration device. The spin column flow through was used as the buffer blank control for SAXS measurements. The samples were snap frozen in liquid nitrogen and stored at -80°C.

### Hydrogen Deuterium Exchange Mass Spectrometry

#### Peptide Identification

Typical HDX-MS peptide identification was performed using a previously described automated liquid handling robot coupled to an Agilent 1260 HPLC and either a Q Exactive (QE) Orbitrap mass spectrometer (Thermo Fisher) [75]. Briefly, 25 μL of 2 μM RORγ2 in HBST buffer (25 mM HEPES pH 7.9, 150 mM NaCl, 1 mM TCEP, 2% glycerol) was mixed with 25 μL of quench buffer (5M urea, 25 mM TCEP, 1.0% TFA) and immediately injected for in-line immobilized pepsin or NepII digestion. Following digestion, peptides were desalted on a trap column (1 × 10 mm C8 HypersilGold, Thermo) and separated on an analytical column (1 × 50 mm C18 HypersilGold, Thermo) using a linear 60-minute linear gradient of 5–40% solvent B (solvent A is 0.3% formic acid and solvent B is 0.3% formic acid 95% acetonitrile). On the QE, master scans were collected at 70,000 K resolution at 400 m/z. Product ion spectra were acquired in data-dependent mode such that the top 10 most abundant ions selected for the product ion analysis by higher-energy collisional dissociation (HCD) between survey scan events. Following MS2 acquisition, the precursor ion was excluded for 16 s. The resulting MS/MS data files were submitted to Mascot (Matrix Science) for peptide identification. Peptides included in the HDX analysis peptide set had a MASCOT score greater than 20 and the MS/MS spectra were verified by manual inspection. The MASCOT search was repeated against a decoy (reverse) sequence and ambiguous identifications were ruled out and not included in the HDX peptide set.

#### Method to Improve Sequence Coverage

Although pepsin digestion provided adequate coverage of the RORγLBD, we found that sequence coverage of the DBD was poor. We hypothesized that this was due to a high charge density of the DBD as pepsin cannot cleave at basic residues [48]. We tried using Nep2 that is capable of cleaving at basic residues [76] and found improved DBD coverage but worsening coverage of the LBD. We hypothesized that HCD fragmentation of high charge state peptides yielded poor fragmentation patterns that were not amenable for database searching. To address these issues, we conceived a strategy to improve coverage using a split charge-state dependent acquisition. To simulate peptide separation during HDX-MS data acquisition, the process was repeated, but the peptides were separated over a 5-minute gradient and only MS1 scans were collected. Like before, quenched protein samples were injected, desalted, and eluted with a 5 min gradient. Instead of analyzing the eluate on the QE (Thermo Fisher), the samples were diverted away and collected in a 1.5 mL Eppendorf tube. The peptide samples were evaporated to dryness using a SPD1010 SpeedVac (thermo) and reconstituted in 25 μL of 0.2 % TFA. 1 μL of each peptide sample (∼250ng) was injected for LC-MS analysis using a Ultimate3000 HPLC system coupled to the Orbitrap Fusion (Thermo). Peptide samples were loaded onto a μPAC trapping column (PharmaFluidics) desalted with loading buffer (2% ACN, 0.1% TFA) flowing at 20 μL/min for 3 minutes. The samples were then separated on a 50 cm μPAC analytical column with a 90-minute gradient (2-30% B in 60 minutes, 30-60% B in 30 minutes; buffer A is 0.1% formic acid and buffer B is 80% acetonitrile and 0.1% formic acid). HPLC eluate was interfaced to on an Orbitrap Fusion Lumos (Thermo) via a nanospray ionization source operating at 2500 V. Master scans were collected on the Orbitrap mass analyzer at 120,000 K resolution at 400 m/z once per second. Candidate precursors above the 1E5 signal threshold were filtered based on monoisotopic mass distribution and then added the dynamic exclusion list for 30 s. Precursor ions were filtered based on charge such that precursors with a charge state whose charges were between +1 and +4 were fragmented by HCD and precursors with charges of +5 to +8 were selected for EThcD. All MS2 spectra were collected in the ion trap mass analyzer operating in normal mode. Raw files containing mixed method fragmentation techniques were processed using Proteome Discoverer (version 2.2.0.338). Only methionine oxidation was included as a dynamic modification during database searches. MS2 spectra were split based on fragmentation technique. HCD and EThcD MS2 spectra were submitted for b/y and c/z ion searches, respectively, with Sequest HT. The databases used for searching contain the target protein and the reverse decoy sequence. All peptide spectral matches were considered for final false discovery rate determination. The PD result file was imported into Skyline (University of Washington) [77] as a spectral library. Transition settings were adjusted such that all MS1 scan resolution reflected the settings of the mass spectrometer HDX-MS conditions (70,000 K resolution in this case). T0 control data (5-minute gradient) collected on the QE was imported as a result file. Precursors were selected based on their fragmentation pattern, mass error (< 10 ppm), isotope distribution (idotp score > 0.9), and whether the flanking cleavage sites abide the “cleavage rules” for pepsin [48] or NEP2 [76]. Peptides were mapped to the primary sequence using a previously described algorithm [78] implemented in HDX-Workbench (Omics Informatics LLC, version 4.2.4). Fast gradient MS1 data was loaded as a 0s on-exchange or T0 control in Workbench. The T0 control was then searched for corresponding ions within a 10-ppm difference from theoretical mass of the identified peptide set sequences. From these settings the amino acid sequence coverage was calculated.

#### HDX-MS analysis

Unless otherwise stated, sample handling and peptide separation were conducted at 4°C. For differential HDX, 50 μL aliquots of 9 μM RORγ:RORE complexes were thawed. Next, 5 μl of sample was diluted into 20 μl D_2_O buffer (25 mM HEPES pH 7.9, 150 mM NaCl, 1 mM TCEP) and incubated for various time points (0, 10, 30, 60, 300, 900, and 3600 s) at 4°C. The deuterium exchange was then slowed by mixing with 25 μL of cold (4°C) 5 M urea, 25 mM TCEP, and 1% TFA. For SRC3 interaction mapping, RORγ2:cRORE was purified and concentrated to 10 μM and treated with 50 μM SR19547 for 10 minutes on ice. The RORγ2:cRORE:SR19547 solution was then diluted with 1:1 with 12 μM SRC3 in protein buffer (50 mM HEPES pH 8.0, 100 mM NaCl, 5 mM DTT, 1M urea, 5% glycerol) or with protein buffer alone. The complexes were incubated for another 30 minutes on ice prior to starting the D_2_O exposure time course. The time course consisted of 5 time points (0, 10, 60, 400, 2820, 14400 s). Quenched samples were immediately injected into the HDX platform. Upon injection, samples were passed through an immobilized pepsin column (2 mm × 2 cm) at 50 μl min^−1^ and the resulting peptides were captured on a 2 mm × 1 cm C8 trap column (Agilent) and desalted for 2.5 minutes. The protease column was housed in a chamber that maintained temperature at 18°C. Peptides were separated across a 2.1 mm × 5 cm C18 column (1.9 μl Hypersil Gold, ThermoFisher) with a linear gradient of 4-40 % CH_3_CN and 0.3 % formic acid, over 5 minutes. Mass spectrometric data were acquired using an Orbitrap mass spectrometer (Q Exactive, Thermo Fisher). The intensity weighted mean m/z centroid value of each peptide envelope was calculated and subsequently converted into a percentage of deuterium incorporation. This was accomplished determining the observed averages of the undeuterated and fully deuterated spectra and using the conventional formula described elsewhere [79]. Corrections for back-exchange were determined empirically using a D_max_ control. Briefly, this sample was generated by mixing 5 μL of protein sample with 20 μL of deuterated buffer and incubated at 37 °C overnight before being queued to for subsequent quenching, and injection.

#### Data Rendering

The HDX data from all overlapping peptides were consolidated to individual amino acid values using a residue averaging approach. Briefly, for each residue, the deuterium incorporation values and peptide lengths from all overlapping peptides were assembled. A weighting function was applied where shorter peptides were weighted more heavily, and longer peptides were weighted less. Each of the weighted deuterium incorporation values were then averaged to produce a single value for each amino acid. The initial two residues of each peptide, as well as prolines, were omitted from the calculations. This approach is similar to that previously described [80]. HDX analyses were performed in triplicate, with single preparations of each purified protein/complex. Statistical significance for the differential HDX data is determined by t-test for each time point and is integrated into the HDX Workbench software [81].

### Crosslinking Mass Spectrometry

#### Crosslinking reaction and sample preparation

RORγ:RORE samples were crosslinked in HBST buffer using disuccinimidyl sulfoxide (DSSO, Cayman chemicals). The reactions were initiated by spiking 1 μL of 75 mM DSSO (Thermo Fisher) in DMSO into a 50 μL solution containing 10 μM protein and mixing. The final molar ratio of crosslinker:protein was 150:1. The reactions were incubated at 25 °C for 45 minutes before being quenched by the addition of Tris pH 8.0 to 50 mM. For SRC3-RORγ crosslinking reactions, 12 μM SRC3 protein samples were dialyzed with HBST (25 mM HEPES pH 7.5 at 25°C, 150 mM NaCl, 1 mM TCEP) for 4 h at 4°C. Dialysis buffer was exchanged every 2 h. The dialyzed SRC3 protein was added 1:1 to RORγ such that the total protein concentration was 10 μM. The DSSO crosslinking reaction was done using a 150 molar fold excess of DSSO to total protein. The crosslinking reaction proceeded for 30 minutes at room temperature. The crosslinking reaction was quenched by the addition of TRIS (pH 8.0) to 50 mM. Each crosslinking reaction was done in triplicate and the reaction replicates were pooled after quenching. The presence of crosslinked protein was confirmed by comparing to the no crosslink negative control samples with SDS-PAGE and Coomassie staining. Non-crosslinked negative controls were generated using the same procedure with vehicle instead of 75 mM DSSO. The remaining DSSO-crosslinked and non-crosslinked samples were then acetone precipitated at - 20 °C overnight and the protein isolated by centrifugation at 16,000 rcf for 5 minutes at 4 °C. After decanting, the pellets were dried for 15 minutes before being resuspended in 12.5 μL of resuspension buffer (50 mM ammonium bicarbonate, 8 M urea, pH 8.0). ProteaseMAX (Promega) was added to 0.02% and the solutions were mixed on an orbital shaker operating at 400 RPM for 5 minutes. After resuspension, 87.5 μL of digestion buffer (50 mM ammonium bicarbonate, pH 8.0) was added. Protein samples were reduced by adding 1 μL of 500 mM DTT and incubating the protein solutions in an orbital shaker operating at 400 RPM and 56°C for 20 minutes. After reduction, 2.7 μL of 550 mM iodoacetamide was added and the solutions were incubated at room temperature in the dark for 15 minutes. Reduced and alkylated protein solutions were digested using trypsin at a ratio of 1:200 (w/w trypsin:protein) and incubating at 37 °C. After 4 h trypsin digestion, reactions were split into 50 μL aliquots and chymotrypsin was added to one aliquot to a final ratio of 1:100 (w/w chymotrypsin:protein). Trypsin and trypsin-chymotrypsin serial digestion reactions were then incubated at room temperature overnight and acidified by addition of TFA to 1 %. Samples were then frozen and stored at -20 °C until analysis.

#### Liquid Chromatography and Mass Spectrometry

Peptide samples were thawed, mixed, and spun down at 16,000 rcf for 5 minutes. 10 μL of peptide samples were loaded in an Ultimate 3000 autosampler (Dionex, ThermoFisher). Approximately 500 ng of each peptide sample was injected in triplicate. Peptides were trapped on a µPAC trapping column (PharmaFluidics) using a load pump operating at 20 μL/min. Load pump buffer contained 2 % acetonitrile and 0.1 % TFA. After a 3-minute desalting period, peptides were then separated using linear gradients (2-30 % Solvent B over 1-60 minutes, 30-95 % solvent B over 60-90 minutes) at 1 μL/min on a 50 cm µPAC C18 column (PharmaFluidics). Gradient solvent A contained 0.1% formic acid and solvent B contained 80% acetonitrile and 0.1% formic acid. liquid chromatography eluate was interfaced to an Orbitrap Fusion Lumos (ThermoFisher) via nano spray ionization source. Crosslinks were identified using a previously described MS2-MS3 method [82]. Master scans of m/z 375-1500 were taken in the orbitrap mass analyzer operating at 60,000 resolution at 400 m/z, 400,000 automatic gain control target. Maximum ion injection time was set to 50 ms and advanced peak detection was enabled. Precursor ions with charge state 4-8 were selected via quadrupole for CID fragmentation at 25 % collision energy and 10 ms reaction time. Fragment ion mass spectrum were taken on the orbitrap mass analyzer operating at 30,000 resolution at 400 m/z, 50,000 automatic gain control target, and maximum injection time was set to 150 ms. Doublet pairs of ions with the targeted mass difference for sulfoxide fragmentation [83] (31.9721 Da) with charge states were selected for HCD fragmentation at 35 % collision energy. MS3 scans were collected in the ion trap operating in ‘rapid’ mode at automatic gain control target and maximum ion injection time set to 20,000 and 200 ms respectively.

#### Data analysis

Crosslinks were identified using the XlinkX algorithm [84] implemented on Proteome Discoverer (version 2.2). Crosslinks were considered for lysine, threonine, serine, and tyrosine residues and the validation strategy was set to ‘simple’ where relaxed and strict false discovery rates were set to 0.05 and 0.01, respectively. During database searches, the target and decoy databases only contained sequences for the proteins that were analyzed, and the maximum missed cleavages was set to 8. Cysteine carbamamidylation and methionine oxidation were considered as a fixed and a dynamic modification, respectively. Mass tolerances for precursor FTMS, fragment FTMS, and fragment ITMS were set to 10 ppm, 20 ppm, and 0.5 Da, respectively. Peak areas for identified crosslinks were quantified using Skyline (version 19.1) using a previously described protocol [85]. Crosslink spectral matches found in Proteome Discoverer were exported and converted to sequence spectrum list format using Excel (Microsoft). Crosslink peak areas were assessed using the MS1 full-scan filtering protocol for peaks within 8 minutes of the crosslink spectral match identification. Peaks areas were assigned to the specified crosslinked peptide identification if the mass error was within 10 ppm of the theoretical mass, the isotope dot product was greater than 0.95, and if the peak was not found in the non-crosslinked negative controls. Pair-wise comparisons were made using ‘MSstats’ package [86] implemented in the Skyline browser to calculate relative fold changes and significance (multiple testing adjusted P-values). Significant changes were defined as log_2_ fold change was less than -1 or greater than 1 and -log_10_ adjusted P-value was greater than 1.3 (P value < 0.05). The results from skyline were exported and are reported in an Excel spreadsheet. Data manipulation, cluster analysis, heatmap plots, and volcano plots were generated with the ‘ggplots2’, ‘gplot’, ‘reshape2’, and ‘factoextra’ packages implemented in R Studio. Crosslinks were mapped to protein sequence using crosslinkviewer [87].

### Small Angle X-ray Scattering

#### Data Collection and Analysis

Dilution series of RORγ:RORE complexes were prepared in 96 well plates with buffer controls (flow through of concentrator device) flanking the dilution series. The highest concentrations ranged from 1.16 to 1.62 mg/mL. The shipping and handling conditions were simulated, and the samples were confirmed to be stable with no detectable aggregates based on analytical size exclusion chromatography. SAXS data collection was done in a high throughput format at the Advanced Light Source beam line 12.4 as previously described [88]. Briefly, SAXS was captured in 0.3 second exposures for 10 s (32 frames total). This data was manually examined and merged using Primus software [89] (part of the ATSAS package of SAXS analysis software [90]). Radius of gyration was determined using the Guinier approximation [91] and the Dmax was determined by distance distribution analysis [92]. The fitted models from pair distance distribution analysis were used as inputs for *ab initio* electron density reconstruction using the DENSS algorithm [53]. Briefly, 50 electron density models were generated and averaged using EMAN2 [93]. The resolution of the electron densities were determined using the gold standard Fourier shell correlation analysis [54].

### Modeling of RORγ:RORE Complexes

Ensembles of models were generated an algorithm that has been previously described [55]. Rigid body models of the LBD were generated from the co-crystal structure (PDB 3L0L) and DBD:RORE were generated by homology modelling. The RORγDBD homology model was generated by Modeler[94] using PDB 1HLZ as a template. Model double stranded DNA oligos were generated using Haddock 3D-DART server [50]. The rigid bodies were connected by the hinge domain that was considered a random coil. These constraints were used to generate 10,000 models and corresponding theoretical SAXS curves. A genetic algorithm that selects combinations of models that collectively fit the experimental SAXS data was run 12 times. This analysis yielded 65 different conformations of RORγ2:RORE. We used these 65 structures as starting models for a simple Rosetta modeling pipeline that rebuilt the hinge region, performed a simple minimization, and then scored the structure using Rosetta energy function REF2015 [95]. Data from XL-MS was incorporated into the scoring function as a flat harmonic Cα distance constrain penalty as previously described [56]. Since the Rosetta pipeline is scalable, we used it to diversify the starting models to 14,417 structures. We scored each of the structures for SAXS agreement using a program called CRYSOL 3.0 [57]. The convergence of the models was assessed by examining the models that were in the top 10% of SAXS χ^2^ and crosslink Cα distance constraint scores. Models and electron densities were visualized using UCSF Chimera[96].

## AKNOWLEGMENTS

We would like to thank the protein production core at Baylor University and, specifically, Professor Dean Edwards and Dr. Yingmin Zhu for providing full length recombinant SRC3 protein. In addition, we would like to thank the beamline scientists at Lawrence-Berkeley National Laboratory for SAXS measurements. This work was conducted at the Advanced Light Source (ALS), a national user facility operated by Lawrence Berkeley National Laboratory on behalf of the Department of Energy, Office of Basic Energy Sciences, through the Integrated Diffraction Analysis Technologies (IDAT) program, supported by DOE Office of Biological and Environmental Research. Additional support comes from the National Institute of Health project ALS-ENABLE (P30 GM124169) and a High-End Instrumentation Grant S10OD018483.

## ACCESSION NUMBERS

Datasets used for exploratory analysis of ChIP-seq datasets from thymocytes, hepatocytes, and TNBC are publicly available in the Gene Expression Omnibus using accessions GSE88916, GSE101115, and GSE126380, respectively. The ChIP- and RNA-seq datasets from Th17 cells can be accessed using accession number GSE40918. HDX- and XL-MS data have been deposited to the ProteomeXchange consortium database[97] and can be accessed using accession identifiers PXD025765 and PXD025728, respectively. The SAXS data for RORγ2:cRORE and RORγ2:vRORE have been deposited to the SASBDB with identifiers SAS3123 and SAS3124, respectively. Scripts used to perform bioinformatic analyses and structural modeling are available from Zenodo using DOI:10.5281/zenodo.4737544.

## REFERENCES

[1] Bookout AL, Jeong Y, Downes M, Yu RT, Evans RM, Mangelsdorf DJ. Anatomical Profiling of Nuclear Receptor Expression Reveals a Hierarchical Transcriptional Network. Cell. 2006;126:789–99.

[2] Medvedev A, Yan ZH, Hirose T, Giguere V, Jetten AM. Cloning of a cDNA encoding the murine orphan receptor RZR/ROR gamma and characterization of its response element. Gene. 1996;181:199–206.

[3] Takeda Y, Kang HS, Freudenberg J, DeGraff LM, Jothi R, Jetten AM. Retinoic acid-related orphan receptor γ (RORγ): a novel participant in the diurnal regulation of hepatic gluconeogenesis and insulin sensitivity. PLoS Genet. 2014;10:e1004331–e.

[4] Meissburger B, Ukropec J, Roeder E, Beaton N, Geiger M, Teupser D, et al. Adipogenesis and insulin sensitivity in obesity are regulated by retinoid-related orphan receptor gamma. EMBO Molecular Medicine. 2011;3:637.

[5] Cai D, Wang J, Gao B, Li J, Wu F, Zou JX, et al. RORγ is a targetable master regulator of cholesterol biosynthesis in a cancer subtype. Nature Communications. 2019;10:4621.

[6] Cai D, Zhang X, Chen H-W. A master regulator of cholesterol biosynthesis constitutes a therapeutic liability of triple negative breast cancer. Molecular & Cellular Oncology. 2020;7:1701362.

[7] Wang J, Zou JX, Xue X, Cai D, Zhang Y, Duan Z, et al. ROR-[gamma] drives androgen receptor expression and represents a therapeutic target in castration-resistant prostate cancer. Nat Med. 2016;22:488–96.

[8] Lytle NK, Ferguson LP, Rajbhandari N, Gilroy K, Fox RG, Deshpande A, et al. A Multiscale Map of the Stem Cell State in Pancreatic Adenocarcinoma. Cell. 2019;177:572-86.e22.

[9] Ivanov II, McKenzie BS, Zhou L, Tadokoro CE, Lepelley A, Lafaille JJ, et al. The Orphan Nuclear Receptor RORγt Directs the Differentiation Program of Proinflammatory IL-17+ T Helper Cells. Cell. 2006;126:1121–33.

[10] Codarri L, Gyülvészi G, Tosevski V, Hesske L, Fontana A, Magnenat L, et al. RORγt drives production of the cytokine GM-CSF in helper T cells, which is essential for the effector phase of autoimmune neuroinflammation. Nature Immunology. 2011;12:560–7.

[11] Ye P, Garvey PB, Zhang P, Nelson S, Bagby G, Summer WR, et al. Interleukin-17 and Lung Host Defense againstKlebsiella pneumoniae Infection. American Journal of Respiratory Cell and Molecular Biology. 2001;25:335–40.

[12] Eberl G. RORγt, a multitask nuclear receptor at mucosal surfaces. Mucosal Immunology. 2017;10:27–34.

[13] Jetten AM, Cook DN. (Inverse) Agonists of Retinoic Acid–Related Orphan Receptor γ: Regulation of Immune Responses, Inflammation, and Autoimmune Disease. Annual Review of Pharmacology and Toxicology. 2020;60:371–90.

[14] Okada S, Markle JG, Deenick EK, Mele F, Averbuch D, Lagos M, et al. Impairment of immunity to *Candida* and *Mycobacterium* in humans with bi-allelic *RORC* mutations. Science. 2015;349:606.

[15] Sun Z, Unutmaz D, Zou Y-R, Sunshine MJ, Pierani A, Brenner-Morton S, et al. Requirement for RORγ in Thymocyte Survival and Lymphoid Organ Development. Science. 2000;288:2369.

[16] Kurebayashi S, Ueda E, Sakaue M, Patel DD, Medvedev A, Zhang F, et al. Retinoid-related orphan receptor γ (RORγ) is essential for lymphoid organogenesis and controls apoptosis during thymopoiesis. Proceedings of the National Academy of Sciences. 2000;97:10132.

[17] Solt LA, Banerjee S, Campbell S, Kamenecka TM, Burris TP. ROR Inverse Agonist Suppresses Insulitis and Prevents Hyperglycemia in a Mouse Model of Type 1 Diabetes. Endocrinology. 2015;156:869–81.

[18] Solt LA, Kumar N, Nuhant P, Wang Y, Lauer JL, Liu J, et al. Suppression of T(H)17 Differentiation and Autoimmunity by a Synthetic ROR Ligand. Nature. 2011;472:491–4.

[19] Chang MR, Lyda B, Kamenecka TM, Griffin PR. Pharmacologic Repression of Retinoic Acid Receptor– Related Orphan Nuclear Receptor γ Is Therapeutic in the Collagen-Induced Arthritis Experimental Model. Arthritis & Rheumatology. 2014;66:579–88.

[20] Liljevald M, Rehnberg M, Söderberg M, Ramnegård M, Börjesson J, Luciani D, et al. Retinoid-related orphan receptor γ (RORγ) adult induced knockout mice develop lymphoblastic lymphoma. Autoimmunity Reviews. 2016;15:1062–70.

[21] Guntermann C, Piaia A, Hamel M-L, Theil D, Rubic-Schneider T, del Rio-Espinola A, et al. Retinoic-acid-orphan-receptor-C inhibition suppresses Th17 cells and induces thymic aberrations. JCI Insight. 2017;2.

[22] Guo Y, MacIsaac KD, Chen Y, Miller RJ, Jain R, Joyce-Shaikh B, et al. Inhibition of RORγT Skews TCRα Gene Rearrangement and Limits T Cell Repertoire Diversity. Cell Reports. 2016;17:3206–18.

[23] Sun N, Guo H, Wang Y. Retinoic acid receptor-related orphan receptor gamma-t (RORγt) inhibitors in clinical development for the treatment of autoimmune diseases: a patent review (2016-present). Expert Opinion on Therapeutic Patents. 2019;29:663–74.

[24] Santori Fabio R, Huang P, van de Pavert Serge A, Douglass Eugene F, Leaver David J, Haubrich Brad A, et al. Identification of Natural RORγ Ligands that Regulate the Development of Lymphoid Cells. Cell Metabolism. 2015;21:286–98.

[25] Hu X, Wang Y, Hao L-Y, Liu X, Lesch CA, Sanchez BM, et al. Sterol metabolism controls TH17 differentiation by generating endogenous RORγ agonists. Nat Chem Biol. 2015;11:141–7.

[26] Jin L, Martynowski D, Zheng S, Wada T, Xie W, Li Y. Structural Basis for Hydroxycholesterols as Natural Ligands of Orphan Nuclear Receptor RORγ. Molecular Endocrinology. 2010;24:923–9.

[27] Strutzenberg TS, Garcia-Ordonez RD, Novick SJ, Park H, Chang MR, Doebellin C, et al. HDX-MS reveals structural determinants for RORγ hyperactivation by synthetic agonists. eLife. 2019;8:e47172.

[28] Ciofani M, Madar A, Galan C, Sellars M, Mace K, Pauli F, et al. A Validated Regulatory Network for Th17 Cell Specification. Cell. 2012;151:289–303.

[29] Zhang Y, Papazyan R, Damle M, Fang B, Jager J, Feng D, et al. The hepatic circadian clock fine-tunes the lipogenic response to feeding through RORα/γ. Genes Dev. 2017;31:1202–11.

[30] He Z, Ma J, Wang R, Zhang J, Huang Z, Wang F, et al. Two amino acid mutation disrupts RORγt function in Th17 differentiation but not thymocyte development. Nature immunology. 2017;18:1128–38.

[31] Rutz S, Eidenschenk C, Kiefer JR, Ouyang W. Post-translational regulation of RORγt—A therapeutic target for the modulation of interleukin-17-mediated responses in autoimmune diseases. Cytokine & Growth Factor Reviews. 2016;30:1–17.

[32] Zhao Q, Khorasanizadeh S, Miyoshi Y, Lazar MA, Rastinejad F. Structural Elements of an Orphan Nuclear Receptor–DNA Complex. Molecular Cell. 1998;1:849–61.

[33] Giguere V, Tini M, Flock G, Ong E, Evans RM, Otulakowski G. Isoform specific amino-terminal domains dictate DNA binding properties of RORalpha, a novel family of orphan nuclear receptors. Genes Dev. 1994;8:538–53.

[34] Meinke G, Sigler PB. DNA-binding mechanism of the monomeric orphan nuclear receptor NGFI-B. Nature Structural Biology. 1999;6:471–7.

[35] Wilson TE, Fahrner TJ, Milbrandt J. The orphan receptors NGFI-B and steroidogenic factor 1 establish monomer binding as a third paradigm of nuclear receptor-DNA interaction. Molecular and Cellular Biology. 1993;13:5794.

[36] Gearhart MD, Holmbeck SMA, Evans RM, Dyson HJ, Wright PE. Monomeric Complex of Human Orphan Estrogen Related Receptor-2 with DNA: A Pseudo-dimer Interface Mediates Extended Half-site Recognition. Journal of Molecular Biology. 2003;327:819–32.

[37] Ueda H, Sun GC, Murata T, Hirose S. A novel DNA-binding motif abuts the zinc finger domain of insect nuclear hormone receptor FTZ-F1 and mouse embryonal long terminal repeat-binding protein. Molecular and Cellular Biology. 1992;12:5667.

[38] Yang XO, Pappu BP, Nurieva R, Akimzhanov A, Kang HS, Chung Y, et al. T Helper 17 Lineage Differentiation Is Programmed by Orphan Nuclear Receptors RORα and RORγ. Immunity. 2008;28:29–39.

[39] Villarino AV, Kanno Y, O’Shea JJ. Mechanisms and consequences of Jak–STAT signaling in the immune system. Nature Immunology. 2017;18:374–84.

[40] Ghoreschi K, Laurence A, Yang X-P, Tato CM, McGeachy MJ, Konkel JE, et al. Generation of pathogenic TH17 cells in the absence of TGF-β signalling. Nature. 2010;467:967–71.

[41] Lee J-Y, Hall JA, Kroehling L, Wu L, Najar T, Nguyen HH, et al. Serum Amyloid A Proteins Induce Pathogenic Th17 Cells and Promote Inflammatory Disease. Cell. 2020;180:79-91.e16.

[42] Lee Y, Awasthi A, Yosef N, Quintana FJ, Xiao S, Peters A, et al. Induction and molecular signature of pathogenic TH17 cells. Nature Immunology. 2012;13:991–9.

[43] Englander JJ, Del Mar C, Li W, Englander SW, Kim JS, Stranz DD, et al. Protein structure change studied by hydrogen-deuterium exchange, functional labeling, and mass spectrometry. Proceedings of the National Academy of Sciences. 2003;100:7057.

[44] de Vera IMS, Zheng J, Novick S, Shang J, Hughes TS, Brust R, et al. Synergistic Regulation of Coregulator/Nuclear Receptor Interaction by Ligand and DNA. Structure. 2017;25:1506-18.e4.

[45] Zhang J, Chalmers MJ, Stayrook KR, Burris LL, Wang Y, Busby SA, et al. DNA binding alters coactivator interaction surfaces of the intact VDR–RXR complex. Nat Struct Mol Biol. 2011;18:556–63.

[46] Zheng J, Chang MR, Stites RE, Wang Y, Bruning JB, Pascal BD, et al. HDX reveals the conformational dynamics of DNA sequence specific VDR co-activator interactions. Nature Communications. 2017;8:923.

[47] Seacrist CD, Kuenze G, Hoffmann RM, Moeller BE, Burke JE, Meiler J, et al. Integrated Structural Modeling of Full-Length LRH-1 Reveals Inter-domain Interactions Contribute to Receptor Structure and Function. Structure. 2020;28:830-46.e9.

[48] Hamuro Y, Coales SJ, Molnar KS, Tuske SJ, Morrow JA. Specificity of immobilized porcine pepsin in H/D exchange compatible conditions. Rapid Communications in Mass Spectrometry. 2008;22:1041–6.

[49] Wang X, Zhang Y, Yang Xuexian O, Nurieva Roza I, Chang Seon H, Ojeda Sandra S, et al. Transcription of Il17 and Il17f Is Controlled by Conserved Noncoding Sequence 2. Immunity. 2012;36:23–31.

[50] van Dijk M, Bonvin AMJJ. 3D-DART: a DNA structure modelling server. Nucleic Acids Research. 2009;37:W235–W9.

[51] Lloyd SP. Least squares quantization in PCM. IEEE Trans Inf Theory. 1982;28:129–36.

[52] Rochel N, Ciesielski F, Godet J, Moman E, Roessle M, Peluso-Iltis C, et al. Common architecture of nuclear receptor heterodimers on DNA direct repeat elements with different spacings. Nature Structural & Molecular Biology. 2011;18:564–70.

[53] Grant TD. Ab initio electron density determination directly from solution scattering data. Nature Methods. 2018;15:191–3.

[54] Galaz-Montoya JG, Hecksel CW, Baldwin PR, Wang E, Weaver SC, Schmid MF, et al. Alignment algorithms and per-particle CTF correction for single particle cryo-electron tomography. Journal of Structural Biology. 2016;194:383–94.

[55] Tria G, Mertens HDT, Kachala M, Svergun DI. Advanced ensemble modelling of flexible macromolecules using X-ray solution scattering. IUCrJ. 2015;2:207–17.

[56] Kahraman A, Herzog F, Leitner A, Rosenberger G, Aebersold R, Malmström L. Cross-Link Guided Molecular Modeling with ROSETTA. PLOS ONE. 2013;8:e73411.

[57] Franke D, Petoukhov MV, Konarev PV, Panjkovich A, Tuukkanen A, Mertens HDT, et al. ATSAS 2.8: a comprehensive data analysis suite for small-angle scattering from macromolecular solutions. Journal of Applied Crystallography. 2017;50:1212–25.

[58] Xie H, Sadim MS, Sun Z. RORγt Recruits Steroid Receptor Coactivators to Ensure Thymocyte Survival. The Journal of Immunology. 2005;175:3800.

[59] Sen S, Wang F, Zhang J, He Z, Ma J, Gwack Y, et al. SRC1 promotes Th17 differentiation by overriding Foxp3 suppression to stimulate RORγt activity in a PKC-θ–dependent manner. Proceedings of the National Academy of Sciences. 2018;115:E458.

[60] He Z, Zhang J, Du Q, Xu J, Gwack Y, Sun Z. SRC3 Is a Cofactor for RORγt in Th17 Differentiation but Not Thymocyte Development. The Journal of Immunology. 2019;202:760.

[61] Wu Q, Nie J, Gao Y, Xu P, Sun Q, Yang J, et al. Reciprocal regulation of RORγt acetylation and function by p300 and HDAC1. Scientific Reports. 2015;5:16355.

[62] Tanaka K, Martinez GJ, Yan X, Long W, Ichiyama K, Chi X, et al. Regulation of Pathogenic T Helper 17 Cell Differentiation by Steroid Receptor Coactivator-3. Cell Reports. 2018;23:2318–29.

[63] Chandra V, Huang P, Hamuro Y, Raghuram S, Wang Y, Burris TP, et al. Structure of the intact PPAR-γ–RXR-α nuclear receptor complex on DNA. Nature. 2008;456:350–6.

[64] Rastinejad F, Wagner T, Zhao Q, Khorasanizadeh S. Structure of the RXR–RAR DNA-binding complex on the retinoic acid response element DR1. The EMBO Journal. 2000;19:1045–54.

[65] Orlov I, Rochel N, Moras D, Klaholz BP. Structure of the full human RXR/VDR nuclear receptor heterodimer complex with its DR3 target DNA. The EMBO journal. 2012;31:291–300.

[66] Yi P, Wang Z, Feng Q, Pintilie Grigore D, Foulds Charles E, Lanz Rainer B, et al. Structure of a Biologically Active Estrogen Receptor-Coactivator Complex on DNA. Molecular Cell. 2015;57:1047–58.

[67] Yu X, Yi P, Hamilton RA, Shen H, Chen M, Foulds CE, et al. Structural Insights of Transcriptionally Active, Full-Length Androgen Receptor Coactivator Complexes. Molecular Cell. 2020;79:812-23.e4.

[68] Lim HW, Kang SG, Ryu JK, Schilling B, Fei M, Lee IS, et al. SIRT1 deacetylates RORγt and enhances Th17 cell generation. Journal of Experimental Medicine. 2015;212:607–17.

[69] Langmead B, Salzberg SL. Fast gapped-read alignment with Bowtie 2. Nature Methods. 2012;9:357–9.

[70] Zhang Y, Liu T, Meyer CA, Eeckhoute J, Johnson DS, Bernstein BE, et al. Model-based Analysis of ChIP-Seq (MACS). Genome Biology. 2008;9:R137.

[71] Heinz S, Benner C, Spann N, Bertolino E, Lin YC, Laslo P, et al. Simple Combinations of Lineage-Determining Transcription Factors Prime cis-Regulatory Elements Required for Macrophage and B Cell Identities. Molecular Cell. 2010;38:576–89.

[72] Yu G, Wang L-G, He Q-Y. ChIPseeker: an R/Bioconductor package for ChIP peak annotation, comparison and visualization. Bioinformatics. 2015;31:2382–3.

[73] Patro R, Duggal G, Love MI, Irizarry RA, Kingsford C. Salmon provides fast and bias-aware quantification of transcript expression. Nature Methods. 2017;14:417–9.

[74] Love MI, Huber W, Anders S. Moderated estimation of fold change and dispersion for RNA-seq data with DESeq2. Genome Biology. 2014;15:550.

[75] Chalmers MJ, Busby SA, Pascal BD, He Y, Hendrickson CL, Marshall AG, et al. Probing Protein Ligand Interactions by Automated Hydrogen/Deuterium Exchange Mass Spectrometry. Analytical Chemistry. 2006;78:1005–14.

[76] Zheng J, Strutzenberg TS, Reich A, Dharmarajan V, Pascal BD, Crynen GC, et al. Comparative Analysis of Cleavage Specificities of Immobilized Porcine Pepsin and Nepenthesin II under Hydrogen/Deuterium Exchange Conditions. Analytical Chemistry. 2020;92:11018–28.

[77] MacLean B, Tomazela DM, Shulman N, Chambers M, Finney GL, Frewen B, et al. Skyline: an open source document editor for creating and analyzing targeted proteomics experiments. Bioinformatics. 2010;26:966–8.

[78] Pascal BD, Willis S, Lauer JL, Landgraf RR, West GM, Marciano D, et al. HDX Workbench: Software for the Analysis of H/D Exchange MS Data. Journal of The American Society for Mass Spectrometry. 2012;23:1512–21.

[79] Zhang Z, Smith DL. Determination of amide hydrogen exchange by mass spectrometry: a new tool for protein structure elucidation. Protein science : a publication of the Protein Society. 1993;2:522–31.

[80] Keppel TR, Weis DD. Mapping residual structure in intrinsically disordered proteins at residue resolution using millisecond hydrogen/deuterium exchange and residue averaging. Journal of the American Society for Mass Spectrometry. 2015;26:547–54.

[81] Pascal BD, Willis S, Lauer JL, Landgraf RR, West GM, Marciano D, et al. HDX workbench: software for the analysis of H/D exchange MS data. Journal of the American Society for Mass Spectrometry. 2012;23:1512–21.

[82] Liu F, Lössl P, Scheltema R, Viner R, Heck AJR. Optimized fragmentation schemes and data analysis strategies for proteome-wide cross-link identification. Nature Communications. 2017;8:15473.

[83] Kao A, Chiu C-l, Vellucci D, Yang Y, Patel VR, Guan S, et al. Development of a Novel Cross-linking Strategy for Fast and Accurate Identification of Cross-linked Peptides of Protein Complexes. Molecular & Cellular Proteomics. 2011;10:M110.002212.

[84] Liu F, Rijkers DTS, Post H, Heck AJR. Proteome-wide profiling of protein assemblies by cross-linking mass spectrometry. Nature Methods. 2015;12:1179–84.

[85] Chen Z, Chen Z, Rappsilber J. A generic solution for quantifying cross-linked peptides using software Skyline. Protocol Exchange. 2018.

[86] Choi M, Chang C-Y, Clough T, Broudy D, Killeen T, MacLean B, et al. MSstats: an R package for statistical analysis of quantitative mass spectrometry-based proteomic experiments. Bioinformatics. 2014;30:2524–6.

[87] Combe CW, Fischer L, Rappsilber J. xiNET: Cross-link Network Maps With Residue Resolution. Molecular & Cellular Proteomics. 2015;14:1137.

[88] Hura GL, Menon AL, Hammel M, Rambo RP, Poole Ii FL, Tsutakawa SE, et al. Robust, high-throughput solution structural analyses by small angle X-ray scattering (SAXS). Nature Methods. 2009;6:606–12.

[89] Konarev PV, Volkov VV, Sokolova AV, Koch MHJ, Svergun DI. PRIMUS: a Windows PC-based system for small-angle scattering data analysis. Journal of Applied Crystallography. 2003;36:1277–82.

[90] Manalastas-Cantos K, Konarev PV, Hajizadeh NR, Kikhney AG, Petoukhov MV, Molodenskiy DS, et al. ATSAS 3.0: expanded functionality and new tools for small-angle scattering data analysis. Journal of Applied Crystallography. 2021;54.

[91] Petoukhov MV, Konarev PV, Kikhney AG, Svergun DI. ATSAS 2.1 - towards automated and web- supported small-angle scattering data analysis. Journal of Applied Crystallography. 2007;40:s223–s8.

[92] Svergun D. Determination of the regularization parameter in indirect-transform methods using perceptual criteria. Journal of Applied Crystallography. 1992;25:495–503.

[93] Tang G, Peng L, Baldwin PR, Mann DS, Jiang W, Rees I, et al. EMAN2: An extensible image processing suite for electron microscopy. Journal of Structural Biology. 2007;157:38–46.

[94] Šali A, Blundell TL. Comparative Protein Modelling by Satisfaction of Spatial Restraints. Journal of Molecular Biology. 1993;234:779–815.

[95] Alford RF, Leaver-Fay A, Jeliazkov JR, O’Meara MJ, DiMaio FP, Park H, et al. The Rosetta All-Atom Energy Function for Macromolecular Modeling and Design. Journal of Chemical Theory and Computation. 2017;13:3031–48.

[96] Pettersen EF, Goddard TD, Huang CC, Couch GS, Greenblatt DM, Meng EC, et al. UCSF Chimera—A visualization system for exploratory research and analysis. Journal of Computational Chemistry. 2004;25:1605–12.

[97] Perez-Riverol Y, Csordas A, Bai J, Bernal-Llinares M, Hewapathirana S, Kundu DJ, et al. The PRIDE database and related tools and resources in 2019: improving support for quantification data. Nucleic Acids Research. 2019;47:D442–D50.

