## Supplemental figures and tables for "Conformational Changes of RORγ During Response Element Recognition and Coregulator Engagement"

**SUPPLEMENTAL FIGURE 1 for FIGURE 1**

**A** All Differentially Expressed Genes

| Name | p-value | Overlap |
|---|---|---|
| T Helper Cell Differentiation | 2.89E-07 | 12.3 % 9/73 |
| IL-23 Signaling Pathway | 1.06E-06 | 15.9 % 7/44 |
| Th17 Activation Pathway | 1.92E-06 | 9.9 % 9/91 |
| PD-1, PD-L1 cancer immunotherapy pathway | 3.52E-04 | 6.6 % 7/106 |
| JAK/Stat Signaling | 4.73E-04 | 7.5 % 6/80 |

**B** Differentially Expressed Genes with Classic-RORE

| Name | p-value | Overlap |
|---|---|---|
| Th17 Activation Pathway | 1.92E-06 | 7.7 % 7/91 |
| T Helper Cell Differentiation | 7.28E-06 | 8.2 % 6/73 |
| Cardiac Hypertrophy Signaling (Enhanced) | 8.37E-05 | 2.4 % 12/496 |
| HMGB1 Signaling | 9.33E-05 | 4.2 % 7/165 |
| IL-23 Signaling Pathway | 1.77E-04 | 9.1 % 4/44 |

**C** Differentially Expressed Genes with Variant-RORE

| Name | p-value | Overlap |
|---|---|---|
| Role of JAK2 in Hormone-like Cytokine Signaling | 1.62E-05 | 8.8 % 3/34 |
| Growth Hormone Signaling | 1.49E-04 | 4.2 % 3/71 |
| JAK/Stat Signaling | 2.12E-04 | 3.8 % 3/80 |
| Prolactin Signaling | 2.20E-04 | 3.7 % 3/81 |
| IL-9 Signaling | 1.05E-03 | 6.1 % 2/33 |

**SUPPLEMENTAL FIGURE 1 for MAIN TEXT FIGURE 1**: Gene set enrichment analysis of RORγ target genes was conducted using Fisher's $\chi^2$ test using the Ingenuity Pathway Analysis software suite. The results for all differentially expressed genes, differentially expressed genes with a classic-RORE on the gene loci, and differentially expressed genes with a variant-RORE on the gene loci are shown in **panels A**, **B**, and **C**, respectively.

### SUPPLEMENTARY TABLE 1 (split for clarity) for FIGURE 2

| Data Set | RORg2:cRORE + SR19547 | RORg2:vRORE + SR19547 | RORg2:SWT1 | RORg2:SWT2 |
|---|---|---|---|---|
| HDX reaction details | 50 mM HEPES, 150 mM NaCl 1 mM TCEP, $pH_{read}$ = 7.90, 80% Deuterium, 4 °C | 50 mM HEPES, 150 mM NaCl 1 mM TCEP, $pH_{read}$ = 7.90, 80% Deuterium, 4 °C | 50 mM HEPES, 150 mM NaCl 1 mM TCEP, $pH_{read}$ = 7.90, 80% Deuterium, 4 °C | 50 mM HEPES, 150 mM NaCl 1 mM TCEP, $pH_{read}$ = 7.90, 80% Deuterium, 4 °C |
| Quench Conditions | 5 M Urea, 50 mM TCEP, 1% TFA | 5 M Urea, 50 mM TCEP, 1% TFA | 5 M Urea, 50 mM TCEP, 1% TFA | 5 M Urea, 50 mM TCEP, 1% TFA |
| Quench Protocol | Mix 25 µL HDX reaction with 25 µL Quench solution | Mix 25 µL HDX reaction with 25 µL Quench solution | Mix 25 µL HDX reaction with 25 µL Quench solution | Mix 25 µL HDX reaction with 25 µL Quench solution |
| HDX time course (min) | 10, 30, 60, 300, 900, 3600 seconds | 0.25, 1, 10, 60, 480, 1440 | 10, 30, 60, 300, 900, 3600 seconds | 0.25, 1, 10, 60, 480, 1440 |
| HDX control samples | Maximally-labeled (T 'infinity') control, 100% H2O (T 'zero') control | Maximally-labeled (T 'infinity') control, 100% H2O (T 'zero') control | Maximally-labeled (T 'infinity') control, 100% H2O (T 'zero') control | Maximally-labeled (T 'infinity') control, 100% H2O (T 'zero') control |
| Back-exchange (mean / IQR) | 83.6% / 23.7% | | | |
| # of Peptides | 77 | 77 | 77 | 77 |
| Sequence coverage | 90.6% | 90.6% | 90.6% | 90.6% |
| Average peptide length / Redundancy | 15.0 / 2.69 | 15.0 / 2.69 | 15.0 / 2.69 | 15.0 / 2.69 |
| Replicates (biological or technical) | 3 (technical) | 3 (technical) | 3 (technical) | 3 (technical) |
| Repeatability (average standard deviation) | 3.89 %D | 4.72 %D | 4.39 %D | 4.33 %D |
| Significant differences definition | Adjusted p-value < 0.05 | | Adjusted p-value < 0.05 | |

| Data Set | RORgLBD | RORg2 |
|---|---|---|
| HDX reaction details | 50 mM HEPES, 150 mM NaCl 1 mM TCEP, $pH_{read}$ = 7.90, 80% Deuterium, 4 °C | 50 mM HEPES, 150 mM NaCl 1 mM TCEP, $pH_{read}$ = 7.90, 80% Deuterium, 4 °C |
| Quench Conditions | 5 M Urea, 50 mM TCEP, 1% TFA | 5 M Urea, 50 mM TCEP, 1% TFA |
| Quench Protocol | Mix 25 µL HDX reaction with 25 µL Quench solution | Mix 25 µL HDX reaction with 25 µL Quench solution |
| HDX time course (min) | 10, 30, 60, 300, 900, 3600 seconds | 0.25, 1, 10, 60, 480, 1440 |
| HDX control samples | Maximally-labeled (T 'infinity') control, 100% H2O (T 'zero') control | Maximally-labeled (T 'infinity') control, 100% H2O (T 'zero') control |
| Back-exchange (mean / IQR) | 83.6% / 23.7% | |
| # of Peptides | 52 | 77 |
| Sequence coverage | 93.3% (of LBD) | 90.6% |
| Average peptide length / Redundancy | 12.3 / 2.69 | 15.0 / 2.69 |
| Replicates (biological or technical) | 3 (technical) | 3 (technical) |
| Repeatability (average standard deviation) | 2.79 %D | 3.79 %D |
| Significant differences definition | Adjusted p-value < 0.05 | |

Supplementary Table reporting key statistics for HDX-MS data presented in FIGURE 2.

### SUPPLEMENTAL FIGURE 2 for FIGURE 4

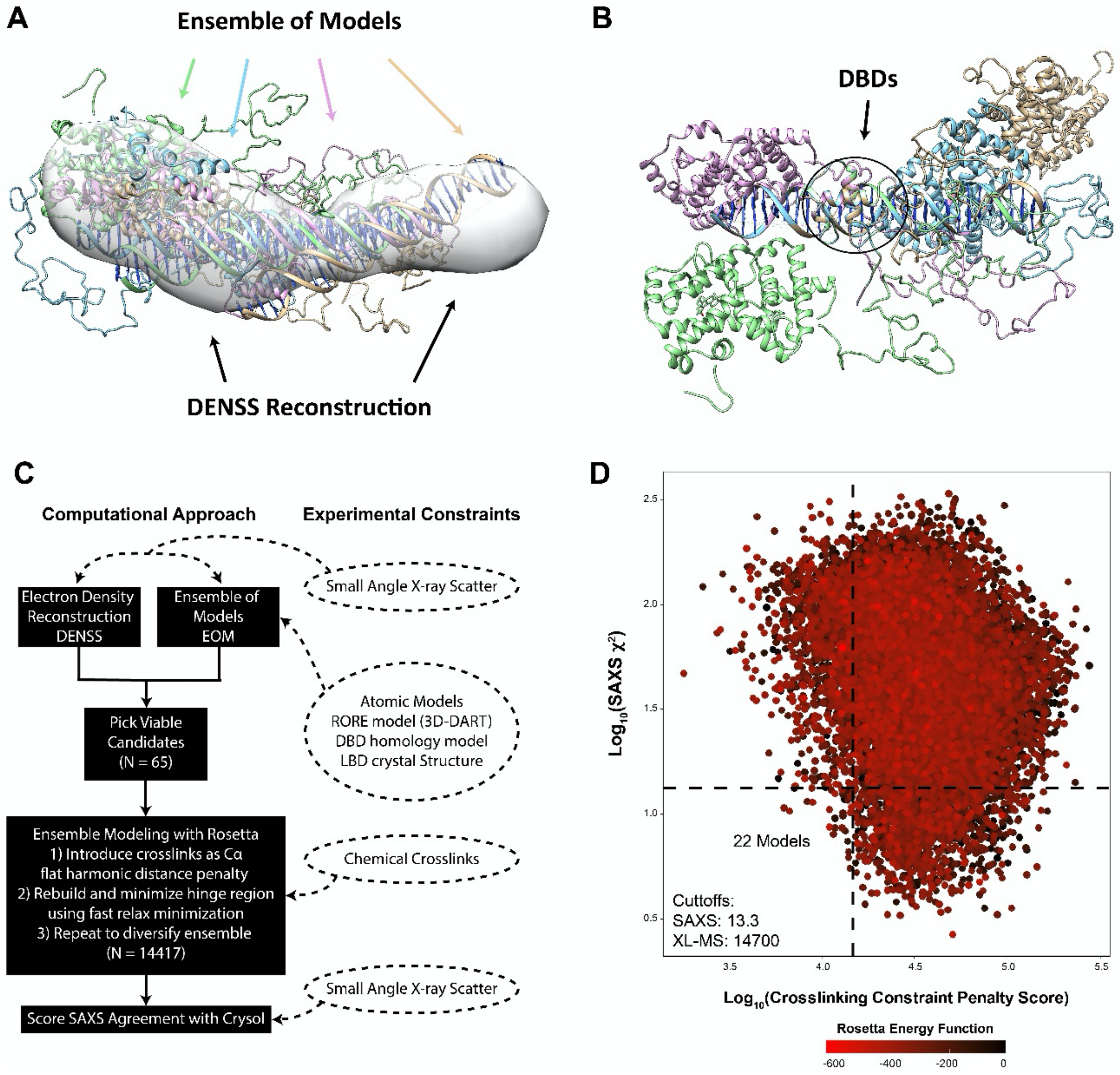


**Supplemental Figure 2 (for main text Figure 4).** An ensemble of models was generated using the EOM algorithm and each model was docked into the electron density reconstruction in **panel A**. The ensemble of models was aligned to the DBD:RORE in **panel B**. This process was repeated until we had 65 starting models that were used as starting structures in a computational modeling approach shown in **panel C**. The distribution of scores is gaussian after $\log_{10}$ transformation of SAXS $\chi^2$ and crosslinking penalty score, as shown in **panel D**. The top scoring models were defined to be models that were in the top 10% for both crosslinking penalty and SAXS $\chi^2$. There were 22 models that satisfied this criterion.

### SUPPLEMENTAL TABLE 2 for FIGURE 5

| Data Set | RORg2:cRORE + SR19547 + Buffer | RORg2:vRORE + SR19547 + SRC3 |
|---|---|---|
| HDX reaction details | 50 mM HEPES, 150 mM NaCl 1 mM TCEP, $pH_{read}$ = 7.90, 80% Deuterium, 4 °C | 50 mM HEPES, 150 mM NaCl 1 mM TCEP, $pH_{read}$ = 7.90, 80% Deuterium, 4 °C |
| Quench Conditions | 5 M Urea, 50 mM TCEP, 1% TFA | 5 M Urea, 50 mM TCEP, 1% TFA |
| Quench Protocol | Mix 25 µL HDX reaction with 25 µL Quench solution | Mix 25 µL HDX reaction with 25 µL Quench solution |
| HDX time course (min) | 10, 60, 400, 2820, 16200 seconds | 10, 60, 400, 2820, 16200 seconds |
| HDX control samples | Maximally-labeled (T 'infinity') control, 100% $H_2O$ (T 'zero') control | Maximally-labeled (T 'infinity') control, 100% $H_2O$ (T 'zero') control |
| Back-exchange (mean / IQR) | 83.6% / 23.7% | |
| # of Peptides | 72 | 72 |
| Sequence coverage | 82.9% | 82.9% |
| Average peptide length / Redundancy | 14.8 / 2.00 | 14.8 / 2.00 |
| Replicates (biological or technical) | 3 (technical) | 3 (technical) |
| Repeatability (average standard deviation) | 2.84 %D | 2.94 %D |
| Significant differences definition | Adjusted p-value < 0.05 | |

Supplementary Table reporting key statistics for HDX-MS data presented in FIGURE 5.
